# Linking acetylated α-Tubulin redistribution to α-Synuclein pathology in brain of Parkinson’s disease patients

**DOI:** 10.1101/2022.12.29.522226

**Authors:** Samanta Mazzetti, Federica Giampietro, Huseyin Berkcan Isilgan, Alessandra Maria Calogero, Gloria Gagliardi, Chiara Rolando, Francesca Cantele, Miriam Ascagni, Manuela Bramerio, Giorgio Giaccone, Ioannis Ugo Isaias, Gianni Pezzoli, Graziella Cappelletti

**Affiliations:** Department of Biosciences, Università degli Studi di Milano, Milan, Italy; Fondazione Grigioni per il Morbo di Parkinson, Milan, Italy; Department of Chemistry, Università degli Studi di Milano, Milan, Italy; Unitech NOLIMITS, Università degli Studi di Milano, 20133 Milan, Italy; S. C. Divisione Oncologia Falck and S. C. Divisione Anatomia Patologica, Ospedale Niguarda Ca’ Granda, Milan, Italy; Unit of Neuropathology and Neurology, Fondazione IRCCS Istituto Neurologico Carlo Besta, Milan, Italy; Parkinson Institute, ASST G. Pini-CTO, Milan, Milan, Italy; Department of Neurology, University Hospital of Würzburg and the Julius Maximilian University of Würzburg, 97080 Würzburg, Germany; Center of Excellence on Neurodegenerative Diseases, Università degli Studi di Milano, Milan, Italy

**Keywords:** Acetylated α-Tubulin, α-Synuclein, oligomers, Lewy body morphogenesis, tunnelling nanotube, Parkinson’s disease

## Abstract

Highly specialized microtubules in neurons are crucial to the health and disease of the nervous system, and their properties are strictly regulated by different post-translational modifications, including α-Tubulin acetylation. An imbalance in the levels of acetylated α-Tubulin has been reported in experimental models of Parkinson’s disease (PD) whereas pharmacological or genetic modulation that leads to increased acetylated α-Tubulin successfully rescues axonal transport defects and inhibits α-Synuclein aggregation. However, the role of acetylation of α-Tubulin in the human nervous system is largely unknown as most studies are based on *in vitro* evidence.

To capture the complexity of the pathological processes *in vivo*, we analysed *post-mortem* human brain of PD patients and control subjects. In the brain of PD patients at Braak stage 6, we found a redistribution of acetylated α-Tubulin, which accumulates in the neuronal cell bodies in subcortical structures but not in the cerebral cortex, and decreases in the axonal compartment, both in the central and peripheral nervous system. High-resolution and 3D reconstruction analysis linked acetylated α-Tubulin redistribution to α-Synuclein oligomerization, leading us to propose a model for Lewy body (LB) morphogenesis. Finally, for the first time in *post-mortem* human brain, we observed threadlike structures, resembling tunnelling nanotubes that contain α-Synuclein oligomers and are associated with acetylated α-Tubulin enriched neurons.

In conclusion, we disclose a novel aspect of LB morphogenesis, indicating the role of acetylated α-Tubulin in PD, that may provide clues to design novel therapeutic interventions.

## 1 Introduction

Cytoskeleton is a pivotal structural component in neurons, where it regulates the architecture and also orchestrates many intracellular events ^1^. Neuronal microtubules are cytoskeletal filaments that are essential for the health of the nervous system not only during neurodevelopment but also in mature neurons as they enable a high degree of morphological and functional complexity, regulate the trafficking of molecules and organelles, and control synaptic plasticity and dendritic spine structure ^2–4^. In the last two decades, microtubule imbalance has emerged as a player in diseases of the nervous system. Different mutations in tubulin genes are linked to a set of neurological disorders, mainly featured by cortical malformation ^5^. Moreover, loss of microtubules or changes in their stability and dynamics may contribute to neurodegeneration as happens in Amyotrophic Lateral Sclerosis, Alzheimer’s disease and Parkinson’s disease (PD) ^6–10^.

Neuronal microtubules are finely regulated by the “tubulin code” that is the result of different α- and ?-tubulin isoforms and a plethora of post-translation modifications (PTMs), which controls both the structure and specific functions of each microtubule ^11–13^. Among these PTMs, acetylation of tubulin has been widely investigated. Lysine 40 acetylation of α-tubulin, found for the first time in Chlamydomonas flagella ^14^, is fundamental for neuronal architecture during development ^15–17^, and is largely present in the axon initial segment and in the axon in mature neurons ^18^. Recently, it has been demonstrated that acetylated α-tubulin enhances microtubule flexibility making them more resilient, protecting them against mechanical aging and stresses ^19^,^20^. In addition, many cellular processes rely on acetylated α-tubulin, including cell migration ^21^, autophagy ^22^, intracellular trafficking ^23^ and cell adhesion ^24^.

The defective regulation of acetylated α-tubulin has been linked to PD ^25–27^. Studies with 1-methyl-4-phenyl-1,2,3,6-tetrahydropyridine (MPTP), and its toxic metabolite 1-methyl-4-phenylpiridinium (MPP+) have shown that the imbalance of acetylated α-tubulin drives cell degeneration. In fact, MPP+ increases the fraction of acetylated α-tubulin and induces defects in axonal transport and mitochondrial damage in PC12 cells ^28^ and in murine primary dopaminergic cultures ^29^. Furthermore, mice treated with MPTP showed an early imbalance of acetylated α-tubulin in nigrostriatal dopaminergic fibers. This is rescued by Epothilone D, a microtubule stabilizer, which attenuates nigrostriatal degeneration showing that microtubule stabilization is neuroprotective ^30^. In addition, recent biochemical data have shown a decrease in acetylated α-tubulin in the *substantia nigra* and no alteration in the cerebral cortex of *post-mortem* human brain obtained from PD patients compared to controls ^31^. PD is the most common neurodegenerative movement disorder ^32,33^, and is characterized by a set of motor and non-motor symptoms and by α-Synuclein aggregation into Lewy bodies (LBs) and Lewy neurites ^34–36^. α-Synuclein aggregation implies the formation of different pathological species including oligomers and fibrils ^37^. α-Synuclein cell-to-cell spreading has been indicated as the route of disease propagation ^38,39^. Studies in cellular models revealed that the transfer can occur thought tunneling nanotubes, exosomes and endocytosis ^40–42^. α-Synuclein aggregation has been linked to the impairment of different cellular processes including regulation of lysosomal and proteasomal functions, mitochondrial activity, biological membrane and synaptic activity, cytoskeleton assembly and functions ^32,33^. Some data in human brain have pointed to α-Synuclein aggregation as a process characterized by different stages, starting with α-Synuclein that accumulates diffusely in neuronal cytoplasm. It then forms a shapeless and irregular aggregate with moderate α-Synuclein staining, and later acquires a well-defined round shape with discrete staining for α-Synuclein in the so-called pale body stage. Finally, it forms an aggregate with a central core and a ring-shaped surrounding halo positive for α-Synuclein that is a typical LB ^43–45^. Interestingly, thanks also to advanced electron microscopy, LBs are defined as crowded aggregates, which in addition to proteins, also contain lipid membranes, vesicular structures, and dysmorphic organelles such as mitochondria and lysosomes ^46^. Very recently, Moors and co-workers successfully used super-resolution microscopy to study the subcellular arrangement of α-Synuclein in PD human brain and demonstrated the presence of an onion skin-type LB containing a well-organized cytoskeletal framework associated with Ser-129-phosphorylated (Ser129 P) α-Synuclein ^47^. Neurofilament and Ēl-Tubulin are distributed mainly in the peripheral part of this structure suggesting that such an organization is important in LB formation ^47^. Additionally, Lewy neurites display cytoskeletal abnormalities due to the possible disruption in the neurofilament network ^46^.

Increasing evidence suggests that α-Synuclein can interact with the microtubular cytoskeleton ^48–50^. Initially, α-Synuclein was found to co-immunoprecipitate with α-tubulin from zebra finch and murine forebrain extracts ^51^. In addition, Alim and colleagues not only confirmed this interaction by demonstrating that α-tubulin co-precipitates with α-Synuclein in the cytoplasm of rat brains ^52^, but also showed that α-Synuclein impacts on microtubule assembly and polymerization *in vitro*^53^. In recent years, many studies have further supported this interaction based on *in vitro* and in cultured cell assays ^50^,^54^, and also on Proximity Ligation Assay (PLA) in mouse brains and in *post-mortem* human brains ^55^. Nevertheless, this interplay seems to be altered in pathological conditions ^49^. Mutated forms of α-Synuclein impact negatively on microtubule assembly *in vitro* and in neuronal cell systems ^50,53,56^ and α-Synuclein overexpression is associated with disruption of the microtubule network causing impairment of axonal transport and neurite degeneration ^57^. As regards human tissues, very few and incomplete data are available. It has been proven that α-Tubulin ^52^ as well as HDAC6, the main α-Tubulin deacetylase, and its phosphorylated active form ^58^ are localised in LBs. However, the mechanism that links α-Tubulin and α-Synuclein during the formation of aggregates warrants further investigation.

Our study shows that acetylated α-tubulin is strongly redistributed in PD brains. First, we identified an accumulation of acetylated α-Tubulin in neuronal cell bodies and a decrease in axons both in the central and peripheral nervous system. We also correlated for the first time acetylated α-Tubulin redistribution with LB formation and analysed in depth the link with α-Synuclein oligomers, which are the early and most aggregating species. Interestingly, we highlighted a driving role of acetylated α-Tubulin during LB morphogenesis in PD human brain.

## 2 Methods

### 2.1 Patients, clinical and neuropathological assessment

All the patients were enrolled in the study and followed during the course of their disease by neurologists experienced in movement disorders and dementia at the ASST G. Pini-CTO Parkinson’s Centre in Milan. Written informed consent was obtained from all subjects prior to enrolment. Clinical diagnosis of PD was established according to the UK Brain Bank criteria ^71,72^. *Post-mortem* human brains were collected by the Nervous Tissues Bank (Milan, Italy) and clinical diagnosis was confirmed by neuropathological analysis, according to the current BrainNet Europe Consortium guidelines ^73^ by two experts, GG and MB. *Post-mortem* human brains obtained from patients fulfilling clinical and neuropathological diagnostic criteria for PD (N = 12) and from age- and sex-matched healthy subjects (N = 6) were used. Demographic data, disease duration and disease severity (according to the Hoehn and Yahr staging system ^74^) are shown in Supplementary Table 1. The study procedures were in accordance with the principles outlined in the Declaration of Helsinki and approved by the Ethics Committee of the University of Milan (protocol code 66/21). Skin biopsies were collected from healthy subjects (N = 9) and PD patients (N = 12) by the Parkinson Institute Biobank ^75^ (Supplementary Table 2).

#### 2.1.1 Human brain samples

Human brains were obtained at autopsy and fixed in 10% buffered formalin for at least 21 days. Specimens from bulb, pons, mesencephalon, basal ganglia, entorhinal, cingulate and the frontal cortex of both controls and PD subjects were embedded in paraffin and then cut into 5-μm thick sections using a microtome (MR2258, Histoline) and processed as follows: i) one slice was stained with haematoxylin and eosin to verify the tissue morphology; ii) one slice was stained for α-Synuclein to verify or exclude its accumulation in LBs and to assess its relative staging, and iii) the sections underwent further immunohistochemistry and immunofluorescence assay for comparative localization of the different antigens.

#### 2.1.2 Skin biopsies

Volar forearm skin biopsies were fixed in Zamboni solution for 24h at 4°C and processed as previously described ^65^. Briefly, the paraffin embedded samples were sliced into 3-μm thick serial sections and processed: i) for histology (haematoxylin and eosin to verify the presence of sweat glands); ii) immunohistochemistry to evaluate the presence of α-Synuclein in the synaptic terminals; and iii) immunofluorescence assay for comparative localization of the different antigens.

### 2.2 Immunohistochemistry

After deparaffination and rehydration, brain sections were sequentially incubated with i) 3% H2O2 for 20 min; ii) 1% BSA diluted in 0.01M phosphate saline buffer (PBS) containing 0.1% Triton™ X-100 (BSAT) for 20 min; iii) the primary mouse anti-acetylated α-Tubulin antibody (6-11B-1), overnight at room temperature (RT). Antigen-antibody interaction was visualized using EnVision™ anti-mouse secondary antibody conjugated with HRP (1 h at RT), and 3,3’ diaminobenzidine as a chromogen (DAB; Dako kit).

In order to assess the comparative localization of acetylated α-Tubulin with α-Synuclein, we performed double immunoenzymatic staining using adjacent sections containing *substantia nigra*, nucleus basalis of Meynert and entorhinal cortex. After incubation with 20% acetic acid for 20 min to inactivate endogenous alkaline phosphatase (AP), the samples were incubated with a mix of mouse anti-acetylated α-Tubulin and rabbit anti α-Synuclein antibody (S3062). To visualize double immunostaining, we used: i) anti rabbit ImmPRESS™-AP secondary antibody conjugated with AP and a substrate solution of 1-Naphthyl phosphate disodium salt (1 mg/ml) and Fast Blue B salt (1 mg/ml) dissolved in 0.1 M Tris-HCl (pH 9.2 - 9.4) for α-Synuclein; and ii) EnVision™ anti-mouse and DAB for acetylated α-Tubulin.

Although we used the well-characterized mouse monoclonal 6-11B-1 for the staining of acetylated α-tubulin ^76^, we checked for its specificity on brain samples. First, primary antibody omission was performed. Next, anti-acetylated α-Tubulin antibody pre-adsorbed with an excess of tubulin (antibody: tubulin 1:5) and an immunoenzymatic assay on *post-mortem* human brain samples (Supplementary Fig. 1). The tubulin necessary for these experiments was obtained in our laboratory from a healthy bovine brain following a well-established protocol ^77^, and an additional chromatographic run using a MonoQ GL column (GE Healthcare Life Sciences). Finally, tubulin was eluted in BRB80 buffer with 1 M KCl, 0.1 mM GTP, 0.1 mM PMSF, 0.1 mM DTT and analysed by SDS-PAGE and western blotting to verify the presence of acetylated α-Tubulin.

### 2.3 Immunofluorescence and Proximity Ligation Assay

Tissue sections containing *substantia nigra*, basal ganglia, frontal cortex and skin biopsies were used for classical immunofluorescence. The tissues were incubated with BSAT for 20 min at RT followed by a mixture of antiacetylated α-Tubulin with different primary antibodies (TH, S100², MBP, IBA1, MAP2, Tau, α-Synuclein, PGP 9.5, Phospho-α-Synuclein; see Supplementary Table 3 for details on antibodies) overnight at RT, followed by incubation with highly pre-adsorbed secondary antibodies (see Supplementary Table 3 for details on antibodies) for 2 h at RT in the dark.

We used PLA to detect α-Synuclein oligomers, as previously described ^65^. Briefly, mesencephalic sections were incubated with a mixture containing α-Synuclein S3062-MINUS, α-Synuclein S3062-PLUS probes, and anti-acetylated α-Tubulin antibody in PLA diluent for 2 h at 37°C and overnight at RT. The amplification reaction was performed in serial incubation steps: i) ligase in Duolink^®^ ligation solution for 1 h at 37°C, and ii) polymerase in Duolink^®^ amplification reagent for 2 h at 37°C, to which donkey anti-mouse secondary antibody was added. Nuclei were stained using Hoechst 33342 (10 min RT) and the samples were mounted using Mowiol-DABCO^®^.

### 2.4 Microscopy and 3D reconstruction

The sections were analysed with a Nikon spinning disk confocal microscope using a water-immersion 40x objective. In addition, we used a spinning disk super-resolution by optical pixel reassignment (SoRa) technique using the silicon-immersion 100x objective plus a resolution improvement of 2.8x. Images were analysed with Fiji software (NIH). For 3D visualisation, images were imported into arivis Vision4D^®^ software (Zeiss Company) containing all the z-stacks in their native format. The images were transformed into the 12-pixel format and region of interest (ROI) were selected using the “Transformation gallery > Crop” tool. Images contain 4 colour channels: Hoechst in blue, acetylated α-Tubulin in red, α-Synuclein oligomers in green and total synuclein in white. The “Intensity threshold segmentation” pipeline was used for the blue and the red signals. The PLA signal showed a scattered and pointy distribution, so the “Blob Finder” pipeline was chosen and using “preview”, a suitable range of exposure was selected. As staining was strong and continuous in a compact area, “Machine Learning Segmentation” was chosen as a suitable pipeline for total-α-Synuclein. Finally, the analysis was run from “Run analysis pipeline” and the reconstruction was visualized via 4D view. The analysis was then carried out using a “cell or particle compartmentalization pipeline”, which can find partial or total overlapping conditions between two reconstructed objects and can be applied to any kind of small particles.

### 2.5 Data and statistical analysis

The data were gathered using Fiji software (NIH). In detail, Fiji tools were used to measure areas, cell counter for counting cells in each anatomic region, and JACOP plug-in for co-localization analysis calculating Mander’s coefficient ^78^. Statistical analysis was carried out using GraphPad Prism 8 and (performing) the Mann-Whitney test. A p-value < 0.05 was considered statistically significant.

In the images obtained using arivis Vision4D^®^ software, the signal was transformed into an object. Data collected from the “object table” by exporting “analysis objects” were the volume of all PLA puncta, the volume of all acetylated α-Tubulin positive objects, the volume of the PLA puncta inside acetylated α-Tubulin positive objects, the volume of the PLA puncta intersecting at least 1% acetylated α-Tubulin positive objects, the mean volume, and the total number of each of the previously listed objects. These data were analysed using frequency distribution and linear correlation, obtaining Spearman’s rank correlation coefficients (ρ).

## DATA AVAILABILITY

The datasets generated during and/or analyzed during the current study are available from the corresponding authors on request.

## 3 Results

### 3.1 Neuronal acetylated α-Tubulin rearrangement in the *substantia nigra* of PD patients

We started by analysing acetylated α-Tubulin distribution in the *substantia nigra*, which is the main region linked to the onset of motor symptoms in PD. In controls, we found that acetylated α-Tubulin is mainly present in cellular processes of neuropil, and surrounds tyrosine hydroxylase (TH) positive neurons, whose cell bodies are mostly negative (Fig. 1a-a’’’). In contrast, in PD samples, acetylated α-Tubulin is enriched in the cellular body (Fig. 1b-b’’’). Therefore, our first goal was to investigate whether this differential distribution of acetylated α-Tubulin is exclusive to neurons or could also be ascribed to glial cells, which are emerging players in PD ^59^. We stained mesencephalic brain sections with acetylated α-Tubulin and glial markers: S100 Calcium Binding Protein β (S100β) for astrocytes, ionized calcium binding adaptor molecule 1 (IBA1) for microglia, and myelin basic protein (MBP) for oligodendrocytes (Supplementary Fig. 2). We observed that astrocytes are enlarged and display more ramifications in PD (Supplementary Fig. 2b, b”) than in controls (Supplementary Fig. 2a, a”). However, quantitative analysis did not reveal any significant difference between control subjects and PD patients in either the percentage of astrocytes positive for acetylated α-Tubulin (Supplementary Fig. 2c) or the co-localization between S100β and acetylated α-Tubulin (Mander’s coefficient; Supplementary Fig. 2d). As for microglia, which has been stated to be highly involved in the phagocytosis of damaged neurons ^60^, the IBA1 marker is clearly visible in the cell bodies (Supplementary Fig. 2e, e”, f, f”). Quantitative analysis revealed no significant difference between control subjects and PD patients in the percentage of microglial cell bodies positive for acetylated α-Tubulin (Supplementary Fig. 2g) and the co-localization between IBA1 and acetylated α-Tubulin (Mander’s coefficient; Supplementary Fig. 2h). Finally, using MBP, we found that acetylated α-Tubulin is mainly distributed along the neuropil in the *substantia nigra* in both control subjects (Supplementary Fig. 2a, a”) and PD patients (Supplementary Fig. 2i, j”), while it is not present in oligodendrocytes. Quantitative analysis demonstrated no significant difference between control subjects and PD patients in terms of MBP co-localisation with acetylated α-Tubulin (Mander’s coefficient; Supplementary Fig. 2k). Collectively, these data reveal that in PD, localisation of acetylated α-Tubulin changes exclusively in neurons.

**Fig. 1.**
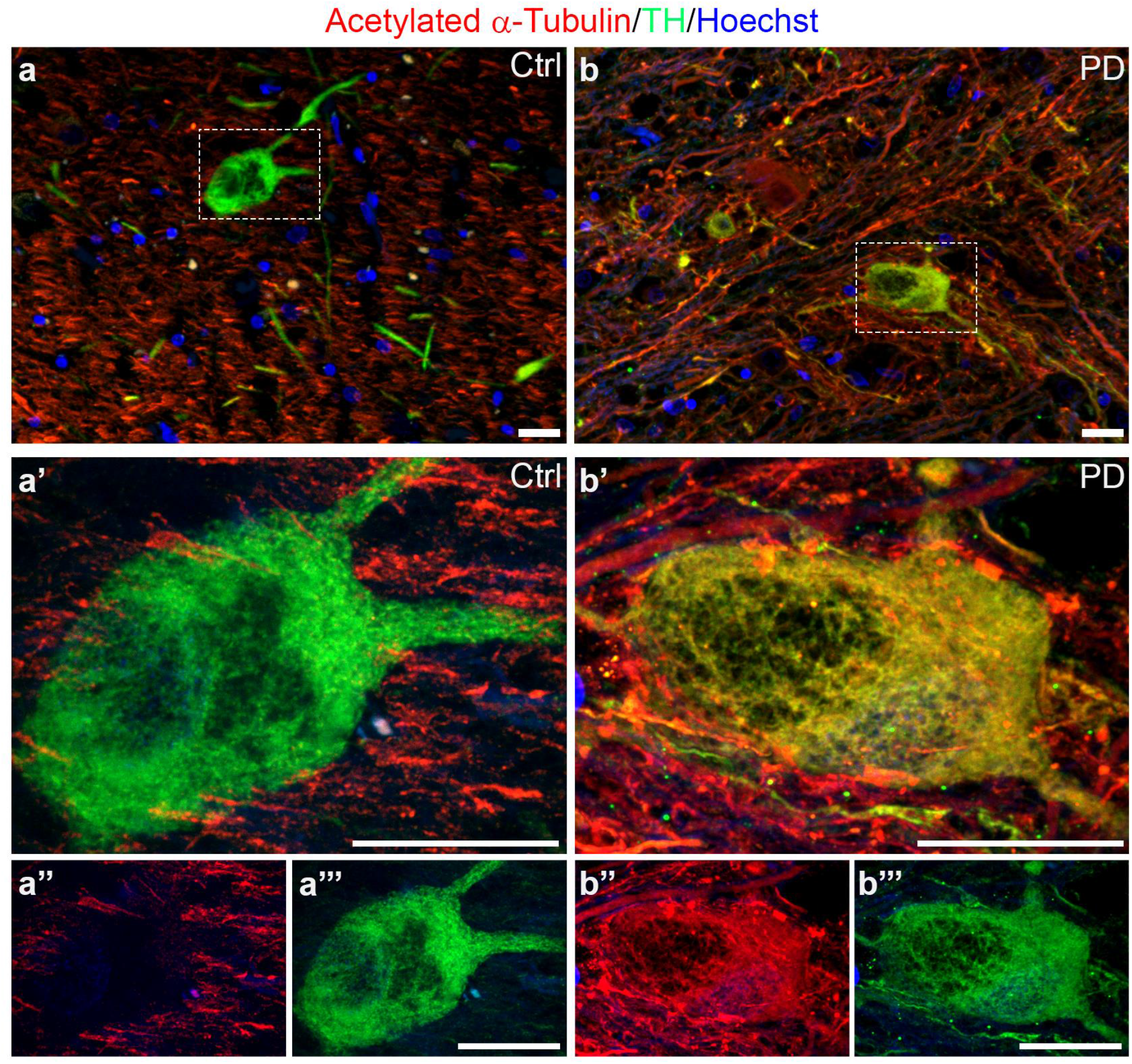
Acetylated α-Tubulin distribution in dopaminergic neurons of *substantia nigra* in *post-mortem* human brain of control subjects (Ctrl) and PD patients (PD). Acetylated α-Tubulin is localised only in neuropil of control samples (**a-a’’’**) while in PD it strongly accumulates within the cell body of dopaminergic neurons (TH positive) containing neuromelanin (**b-b’’’**). Nuclei are counterstained with Hoechst. Scale bar, 20 μm.

### 3.2 Acetylated α-Tubulin mislocalizes in subcortical regions in PD patients

We analysed acetylated α-Tubulin in PD patients, starting from representative subcortical regions to the cerebral cortex ^61^. In controls (Fig. 2a, d, g), acetylated α-Tubulin was mainly localised in the neuropil, as expected, and, rarely, in neuronal cellular bodies in the *substantia nigra, locus coeruleus* and dorsal motor nucleus of vagus, while in PD, neurons with a cell body strongly positive for acetylated α-Tubulin were abundant and greatly increased in all three regions (Fig. 2b, c, e, f, h, i).

**Fig. 2.**
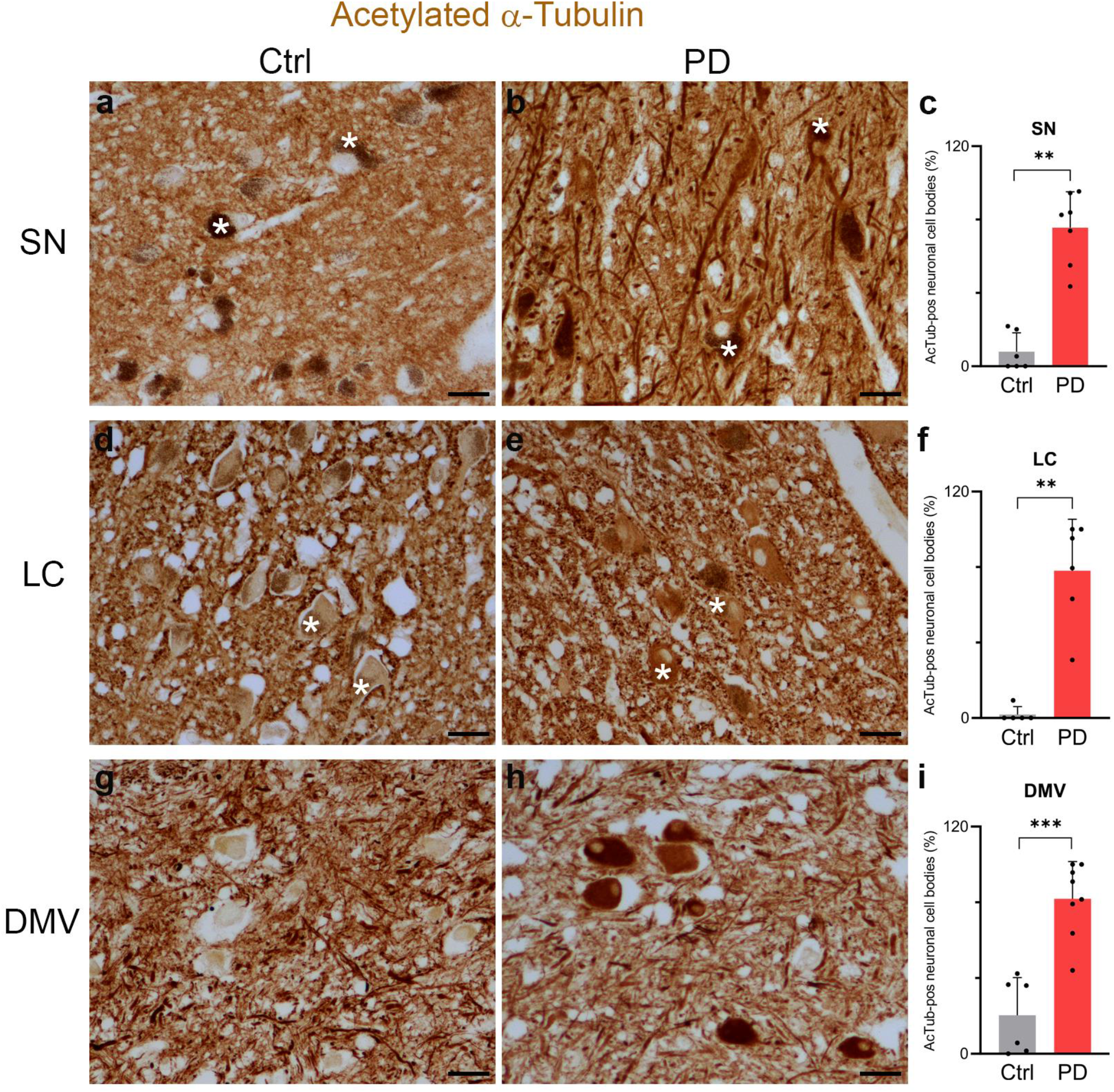
Acetylated α-Tubulin distribution in *post-mortem* human brain. Control samples show acetylated α-Tubulin only in neuropil and in axonal fibres in SN (**a**), LC (**d**) and DMV (**g**). In the same regions of PD samples (**b, e, h**), acetylated α-Tubulin is strongly localised and accumulated in neuronal cytoplasm, and is also present in neuronal processes. The graphs show the percentage of acetylated α-Tubulin positive neuronal cell bodies (**c**: SN Ctrl, N = 6, 263 neurons vs PD, N = 7, 496 neurons; **f**: LC Ctrl, N = 6, 259 neurons vs PD, N = 6, 131 neurons; **i**: DMV Ctrl, N = 6, 334 neurons vs PD, N = 8, 404 neurons). White asterisk: neurons containing neuromelanin. Scale bar, 40 μm. Mann-Whitney test, ** p < 0.01; *** p <0.001. SN: *substantia nigra;* LC: *locus coeruleus;* DMV: dorsal motor nucleus of vagus.

Notably, no differences were observed in the cerebral cortex. Indeed, we investigated the frontal cortex (Supplementary Fig. 3a-e), cingulate cortex (Supplementary Fig. 3f-j) and entorhinal cortex (Supplementary Fig. 3k-o) and found that staining was intense in the apical dendrites (Supplementary Fig. 3a, c, f, h, k, m) and in a limited number of pyramidal neurons (Supplementary Fig. 3b, d, g, i, l, n) in both control and PD samples. However, the non-significant changes in acetylated α-Tubulin could be due to the complexity of the cerebral cortex added to the intrinsic variability of human samples (Supplementary Fig. 3e, j, o). We then investigated in-depth the compartmentalization of acetylated α-Tubulin in cortical neurons. First, we focused on apical dendrites. We double-stained frontal cortex slides with antibodies against acetylated α-Tubulin and MAP2, which is a marker of the somatodendritic compartment. MAP2 was present in the cellular bodies and apical dendrites both in control (Supplementary Fig. 3p, p’’) and PD samples (Supplementary Fig. 3q, q’’). Quantitative analysis revealed no significant difference between control subjects and PD patients in terms of either apical dendrites positive for acetylated α-Tubulin (Supplementary Fig. 3r) or co-localization between MAP2 and acetylated α-Tubulin (Mander’s coefficient; Supplementary Fig. 3s). Furthermore, we investigated the axonal compartment stained for Tau (Supplementary Fig. 4) and found that no differences in the distribution of acetylated α-Tubulin and Tau are detectable in the white matter among controls and PD patients.

Our data disclose the region-specific enrichment of acetylated α-Tubulin in neuronal cell body in PD patients.

### 3.3 Acetylated α-Tubulin is reduced in fibre bundles of putamen

In addition to the cortex, we studied the axonal compartment in other brain regions that are involved in PD, such as the putamen, which plays a role in movement control through direct and indirect circuits. The axonal myelinated bundles, which cross the grey matter, displayed intense staining for acetylated α-Tubulin in control subjects (Fig. 3a), but were weakly stained in PD patients (Fig. 3b). On the contrary, neuron cell bodies lacked acetylated α-Tubulin in controls (Fig. 3a’) but were strongly positive in PD (Fig. 3b’).

**Fig. 3.**
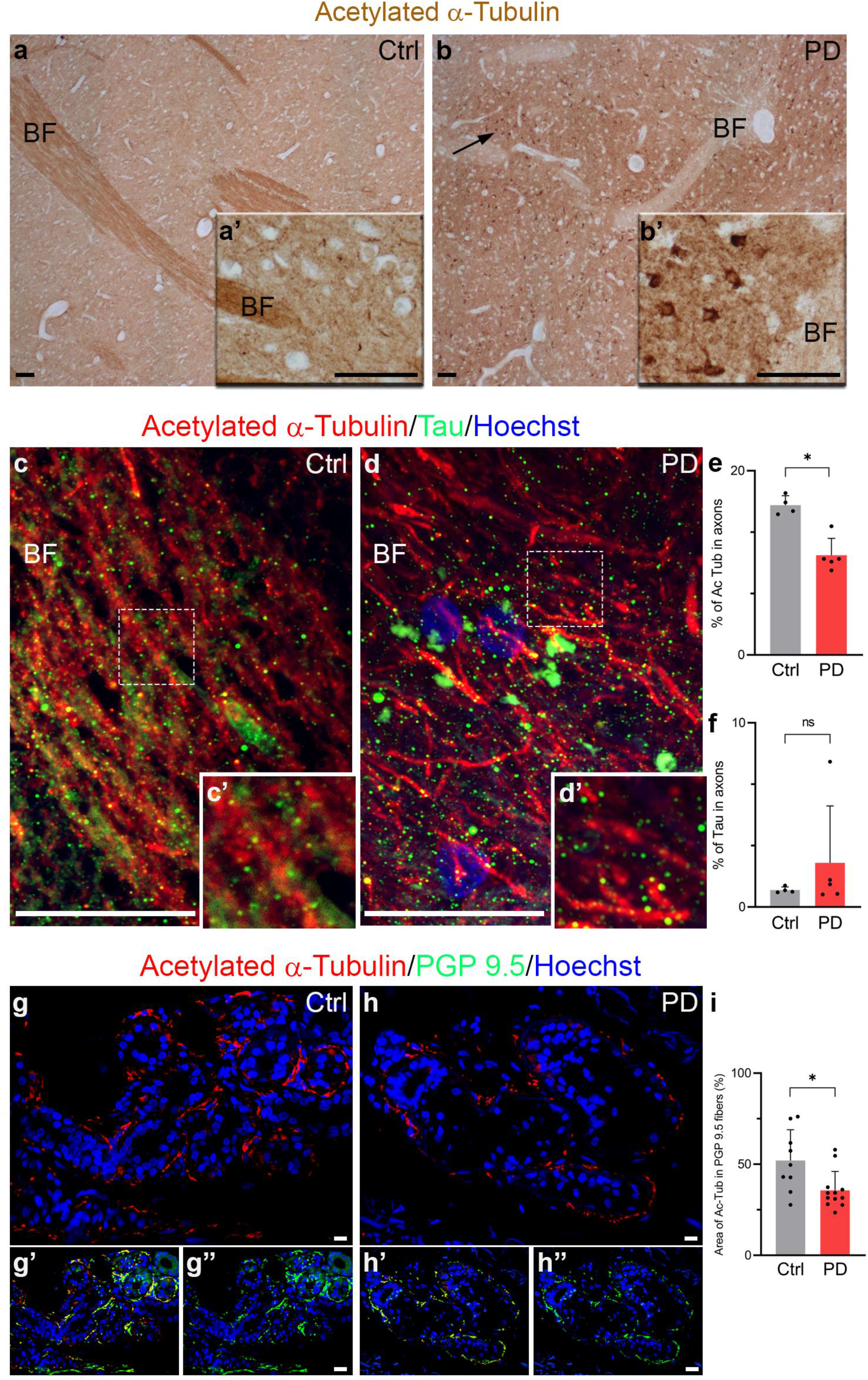
Acetylated α-Tubulin distribution in the fibres of the central and peripheral nervous system. **a-b** Acetylated α-Tubulin is localised in nigro-striatal fibres of putamen but not in neuronal cell bodies in controls (**a-a’**), whereas it is enriched mainly in neuronal cell bodies in PD patients (**b, b’**; black arrow). Scale bar, 100 μm. **c-f** In controls, acetylated α-Tubulin positive axons in putamen run parallel to each other (**c, c’**; dashed square: 2x magnified) while they are less dense in PD samples (**d, d’**; dashed square: 2x magnified). Nuclei are counterstained with Hoechst. The graphs show the percentage of the area covered by acetylated α-Tubulin (**e**; Ctrl, N = 3 vs PD, N = 4) and Tau (**f**; Ctrl, N = 3 vs PD, N = 4) in axons. Scale bar, 20 μm. Mann-Whitney test, * p < 0.05. BF: bundle of fibres. **g-i** Acetylated α-Tubulin localisation within PGP 9.5 positive fibres around sweat glands both in control subjects (**g-g’’**) and PD patients (**h-h’’**). Nuclei are counterstained with Hoechst. The graph (**i** Ctrl, N = 9 vs PD, N = 12) indicates the percentage of the area covered by acetylated α-Tubulin in PGP 9.5 positive fibres. Scale bar, 20 μm. Mann-Whitney test, * p < 0.05.

We used confocal microscopy to perform quantitative analysis. Acetylated α-Tubulin was distributed along the axonal fibres positive for Tau staining (Fig. 3c-d’). Acetylated α-Tubulin was significantly decreased in PD patients (Fig. 3e), while Tau staining did not change (Fig. 3f). This decrease in acetylated α-Tubulin in axons may correspond to the redistribution that we observed in the cellular bodies of neurons in the *substantia nigra, locus coeruleus* and dorsal motor nucleus of vagus as it could reflect the translocation of acetylated α-Tubulin from the axons to the cellular bodies.

### 3.4 Acetylated α-Tubulin is reduced in PNS autonomic fibres in PD patients

Recently, PD has been defined as a multisystem disorder ^62^ which does not only involve the central nervous system but also the peripheral nervous system. Notably, some studies have shown that α-Synuclein aggregation occurs also in the autonomic structure of the skin in PD patients ^63–65^. In particular, PD patients experienced sweating alterations, and we found a significant increase in α-Synuclein oligomers in the autonomic fibres innervating the sweat glands ^65^. This led us to wonder whether the decrease in acetylated α-Tubulin that we saw in the central nervous system could be present also in the autonomic fibres that innervate the sweat glands in skin biopsies. We performed double immunofluorescence for acetylated α-Tubulin and PGP 9.5, which is a marker for fibres. We observed that acetylated α-Tubulin (Fig. 3g-g’, h-h’) localises inside PGP 9.5 positive fibres (Fig. 3g’-g’’, h’-h’’), which surround the sweat glands, both in control and PD samples. Quantitative analysis revealed a significant decrease in acetylated α-Tubulin in the fibres of PD patients (Fig. 3i) that is in line with results obtained in the putamen.

### 3.5 Acetylated α-Tubulin interplay with α-Synuclein aggregation into LB

To highlight the potential role of the redistribution of acetylated α-Tubulin in PD, the relationship between α-Synuclein pathology and acetylated α-Tubulin was investigated, using a double immunoenzymatic method in those regions where LBs are abundant ^61^ and concomitant acetylated α-Tubulin accumulates in the soma: *substantia nigra* (Fig. 4a), nucleus basalis of Meynert (Fig. 4b), and entorhinal cortex (Fig. 4c). In all these areas, four populations of neurons were identified and scored: i) neurons with cell bodies positive for acetylated α-Tubulin; ii) neurons containing LBs; iii) neurons positive for acetylated α-Tubulin and containing LBs; iv) neurons negative for both acetylated α-Tubulin and LBs. Despite the loss of neurons that characterizes the *substantia nigra*, the highest percentage of cells displaying acetylated α-Tubulin accumulation was in this region (Fig. 4a’). Furthermore, in all the areas analysed we found that acetylated α-Tubulin positive cells were approximately 40-70% (Fig. 4a’-c’), while neurons positive for acetylated α-Tubulin and containing LBs were rare (1 – 5 %; Fig 4a’-c’), as confirmed by confocal microscopy (Fig. 4d-e”). Rarely did we observe LBs positive for acetylated α-Tubulin (Fig. 4 e-e”), whereas they were strongly positive for Serl29P α-Synuclein (Supplementary Fig. 5), a marker used for LBs and α-Synuclein pathology ^66^. Taken together, these data suggest that neurons positive for acetylated α-Tubulin and those containing LB tend to be mutually exclusive.

**Fig. 4.**
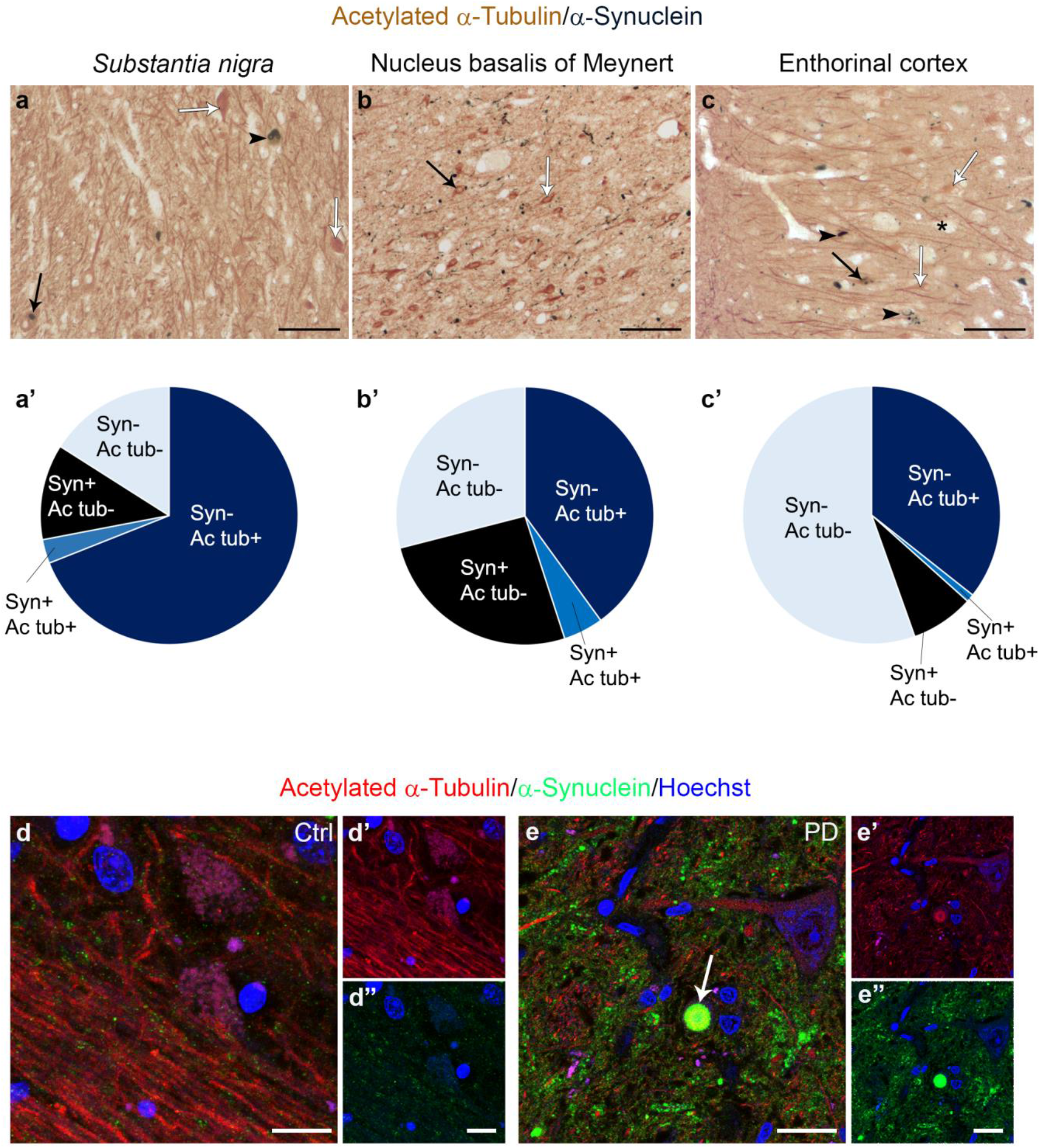
Linking acetylated α-Tubulin redistribution with α-Synuclein pathology in human brain. **a-c’** Four categories of cells are distinguishable in the *substantia nigra* (**a**, N = 3, 574 neurons), nucleus basalis of Meynert (**b**, N = 3, 154 neurons) and entorhinal cortex (**c**, N = 3, 1173 neurons) of PD patients: cells exclusively positive for acetylated α-Tubulin (white arrow), neurons containing α-Synuclein accumulated as Lewy bodies (black arrowheads), cells positive for both acetylated α-Tubulin and α-Synuclein (black arrow) and cells lacking staining (asterisk). Graphs (**a’-c’**) show cell percentages? for each category in the three regions. Scale bar, 100 μm. d/e In controls, acetylated α-Tubulin is present in fibres (**d-d’**) and α-Synuclein shows synaptic staining (**d, d’’**). In PD samples, acetylated α-Tubulin accumulates in neuronal cell bodies and in few α-Synuclein positive aggregates (**e-e’’**, white arrow). Nuclei are counterstained with Hoechst. Scale bar, 25 μm. Ac tub: acetylated α-Tubulin; Syn: α-Synuclein.

Proof that LBs are rarely present in the soma of neurons with increased acetylated α-Tubulin strongly suggests a link between redistribution of acetylated α-Tubulin and the morphogenesis of LBs. We therefore focused on α-Synuclein oligomers ^67^, which are considered the earliest and most toxic species of α-Synuclein during its aggregation process ^37^. Using super-resolution microscopy, we found a staging-like distribution between α-Synuclein and acetylated α-Tubulin. The combination between PLA to detect α-Synuclein oligomers and classical immunofluorescence for acetylated α-Tubulin allowed us to identify different stages that could describe LB morphogenesis (Fig 5): stage 1, redistribution is already present as the cytoplasm of neurons is rich in acetylated α-Tubulin; stage 2, some oligomers are spread in the cytoplasm in which acetylated α-Tubulin is widespread; stage 3, acetylated α-Tubulin starts to accumulate in small aggregates containing oligomers; stage 4, acetylated α-Tubulin is entirely incorporated together with α-Synuclein oligomers inside the aggregate, whose shape is undefined; stage 5, inclusions are roundish with an external ring-shaped structure composed of acetylated α-Tubulin and oligomers, which are also found in the centre of the structure; stage 6, oligomers are localized mainly in the periphery of the aggregate, acetylated α-Tubulin staining is weak while diffuse Hoechst staining is detectable in the core region. All these data highlight for the first time the involvement of acetylated α-Tubulin and α-Synuclein oligomers in the complex process of LB morphogenesis.

**Fig. 5.**
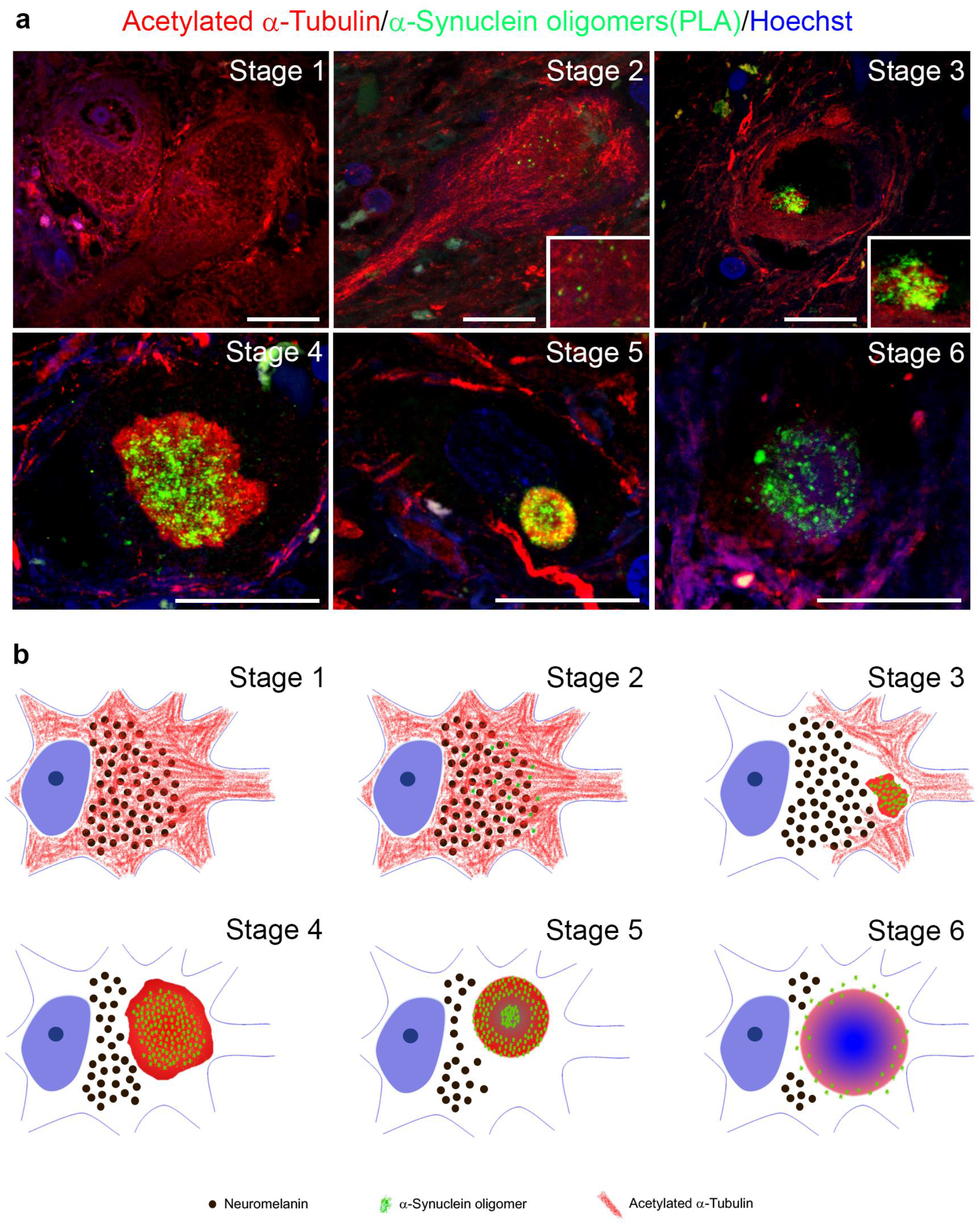
Linking acetylated α-Tubulin redistribution with α-Synuclein oligomerization and Lewy body morphogenesis. **a** Six different stages are distinguishable in *post-mortem* human brain sections of PD patients following PLA assay for detecting α-Synuclein oligomers (green) and immunofluorescence for detecting acetylated α-Tubulin (red). Stage 1 shows the strong presence of acetylated α-Tubulin accumulated in the soma of neurons; in stage 2, acetylated α-Tubulin is still accumulated inside the cell body while some small and spared oligomers appear; in stage 3, acetylated α-Tubulin starts to accumulate into a small aggregate with α-Synuclein oligomers; stage 4 shows acetylated α-Tubulin in a bigger aggregate with an undefined shape with oligomers inside, whereas the cytoplasm is now negative for acetylated α-Tubulin staining; stage 5 shows a ring-shaped aggregate where acetylated α-Tubulin forms an external ring and α-Synuclein oligomers are distributed not only along the ring but also inside it; in stage 6, the ring-shaped aggregate is weakly positive for acetylated α-Tubulin and has few oligomers. Hoechst staining (blue) is detectable in the core of aggregates at stage 6. Scale bar, 20 μm. **b** Schematic model showing the hypothetical sequences of events in LB morphogenesis starting from the redistribution of acetylated α-Tubulin to α-Synuclein oligomerization and aggregation in dopaminergic neurons of PD human brain.

We further investigated the interplay between acetylated α-Tubulin and α-Synuclein oligomers by applying the imaging arivis Vision4D^®^ software on high-resolution images (Fig. 6–7; Supplementary movies). This analytical program guarantees precise and reproducible quantitative morphometric analyses ^68–70^. This is particularly important for visualizing complex structures and their relationships, such as LB morphogenesis and the possible correlation between the acetylated α-Tubulin and α-Synuclein oligomers. Figure 6 reports representative images of aggregates from stage 3 to stage 6 that show 3D reconstructions obtained using a confocal microscope (Fig. 6a, b, c, d) and processed by arivis Vision4D^®^ software (Fig. 6a’-a’’’, b’-b’’’, c’-c’’’, d’-d’’’). These analyses enabled us to explore in detail how α-Synuclein oligomers and acetylated α-Tubulin are distributed inside the different aggregates. Having confirmed the progressive changes in the localization of α-Synuclein oligomers inside the aggregates, we performed further in-depth measurements of PLA puncta in two main types of aggregates: i) undefined aggregates, as observed in stages 3 and 4 (Fig. 7a-b’’), and ii) ring-shaped aggregates, as observed in stages 5 and 6 (Fig. 7d-e’’). To better differentiate the two groups of aggregates, we also investigated total α-Synuclein staining (Fig. 7a, b”, d, e”). The average volume of PLA puncta was lower in undefined aggregates (0.029 ±3.197×10^-6^μm3, n. of PLA puncta = 26712 in 17 aggregates; Supplementary Fig. 6) compared to ring-shaped aggregates (0.045 ±5.534×10^-6^ μm3, n. of PLA puncta =17889 in 15 aggregates; Supplementary Fig. 6), which is in line with progressive α-Synuclein aggregation. Furthermore, we analysed the correlation between the volume of PLA puncta inside acetylated α-Tubulin positive structures and the acetylated α-Tubulin total volume. We found a correlation in both undefined (Fig. 7c; index correlation: 0.77) and ring-shaped (Fig. 7f; index correlation: 0.54) aggregates, supporting the hypothesis that α-Synuclein aggregation is strictly linked to the formation of acetylated α-Tubulin positive structures.

**Fig. 6.**
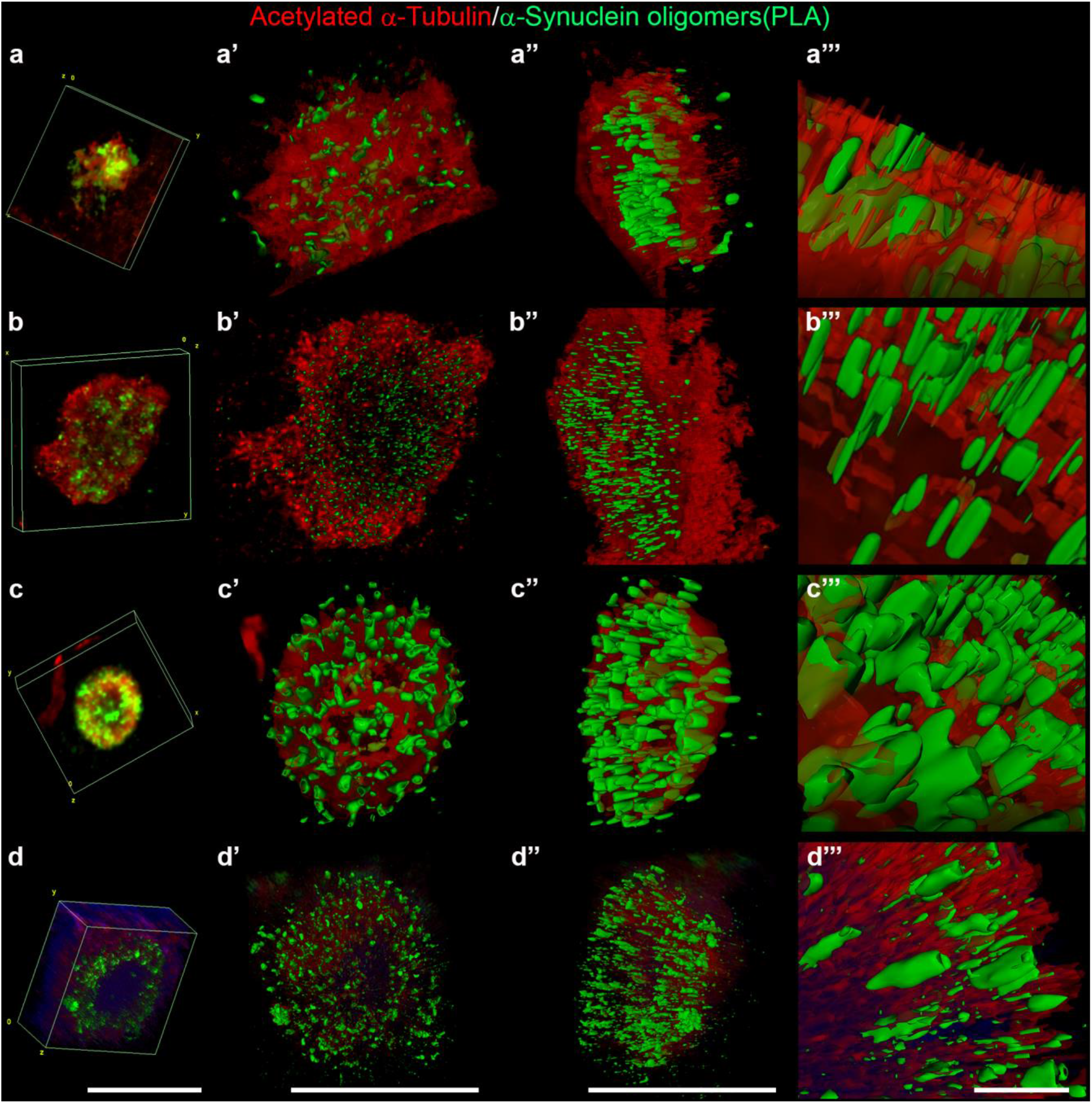
Representative 3D reconstruction of four stages of Lewy body morphogenesis (stage 3-6) obtained with confocal and arivis Vision4D^®^ software. **a-a’’’** stage 3; **b-b’’’** stage 4; **c-c’’’** stage 5; **d-d’’’** stage 6. **a-a’’**, **b-b’’**, **c-c’’**, **d-d’’**: scale bar, 20 μm; **a’’’-d’’’**: scale bar, 5 μm. **a-d**: confocal 3D reconstruction obtained with Fiji software; **a’-d’**: frontal view of 3D reconstruction by arivis Vision4D^®^ software; **a’’-d’’**: lateral view of 3D reconstruction by arivis Vision4D^®^ software; **a’’’-d’’’**: magnifications by arivis Vision4D^®^ software.

**Fig. 7.**
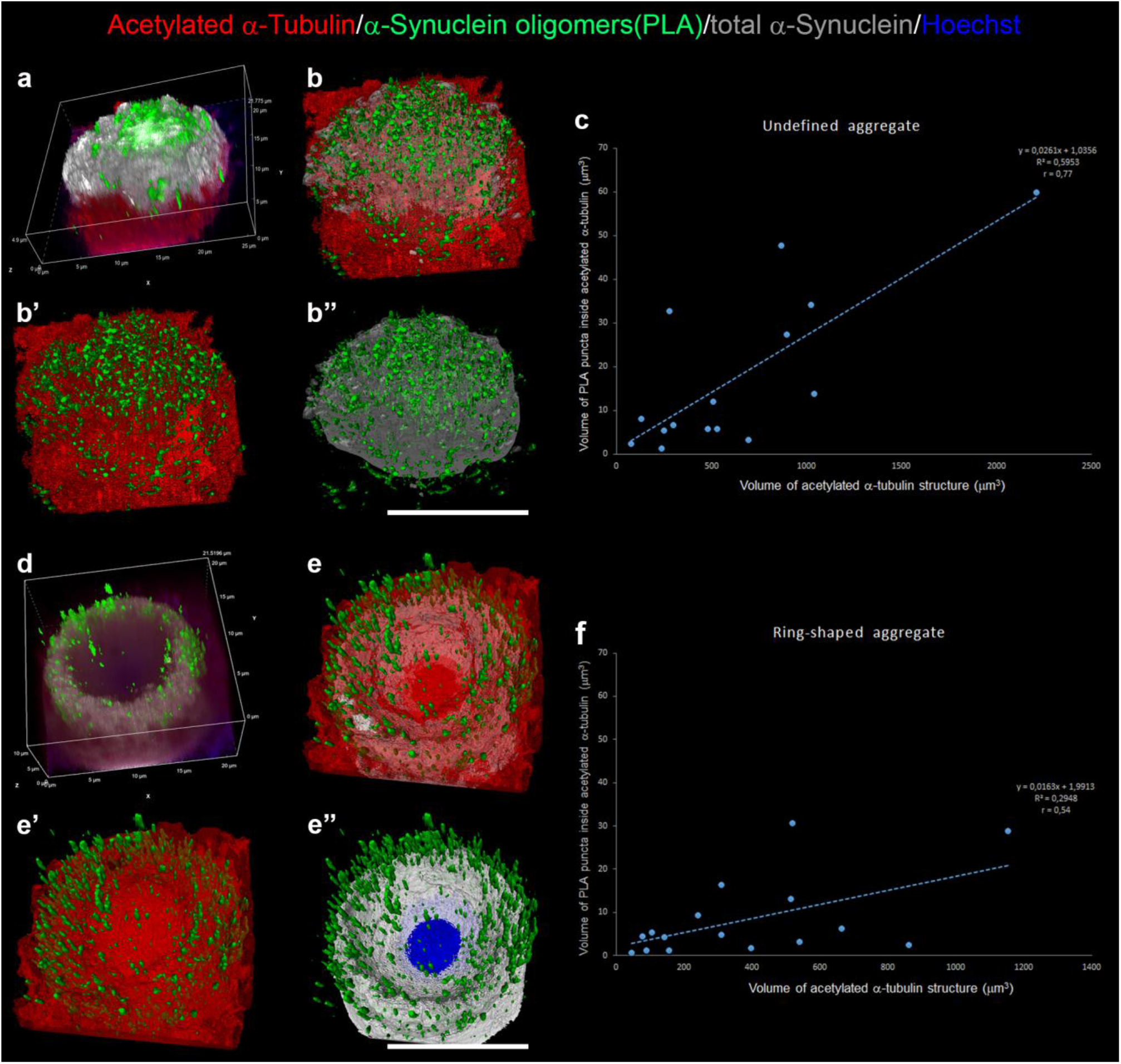
Correlation of α-Synuclein oligomers and acetylated α-Tubulin positive structures between undefined and ring-shaped aggregates. a/b’’ 3D reconstruction obtained with Nis Elements software (**a**) and arivis Vision4D^®^ software (**b-b’’**) of an undefined aggregate. Graph (**c**) shows the linear correlation between the volume of α-Synuclein oligomers and volume of acetylated α-Tubulin positive structures. d/f 3D reconstruction obtained with Nis Elements software(**d**) and arivis Vision4D^®^ software (**e-e’’**) of a ring-shaped aggregate. Graph (**f**) shows the linear correlation between the volume of α-Synuclein oligomers and the volume of acetylated α-Tubulin positive structures. Scale bar, 10 μm.

Finally, the extensive analysis of brain sections that we stained for α-Synuclein oligomers and acetylated α-Tubulin revealed a structure, that resembles the so-called tunnelling nanotube, previously described mainly *in vitro* in cellular models ^38,40,41^, but never found in human brain. Indeed, for the first time, we observed 6 threadlike structures in the *substantia nigra* detected by the presence of α-Synuclein oligomers in different patients. In detail, the tunnelling nanotubes are located between a neuron containing an acetylated α-Tubulin positive structure and another neuron (Fig. 8) or, alternatively, vessels and glial cells (Supplementary Fig. 7). 3D reconstruction enables us to better visualize the distribution of α-Synuclein oligomers in the sample (Fig. 8 b-b”). The structures we identified could indicate the spreading of oligomers, confirming the diffusive properties of this species also in human brain.

**Fig. 8.**
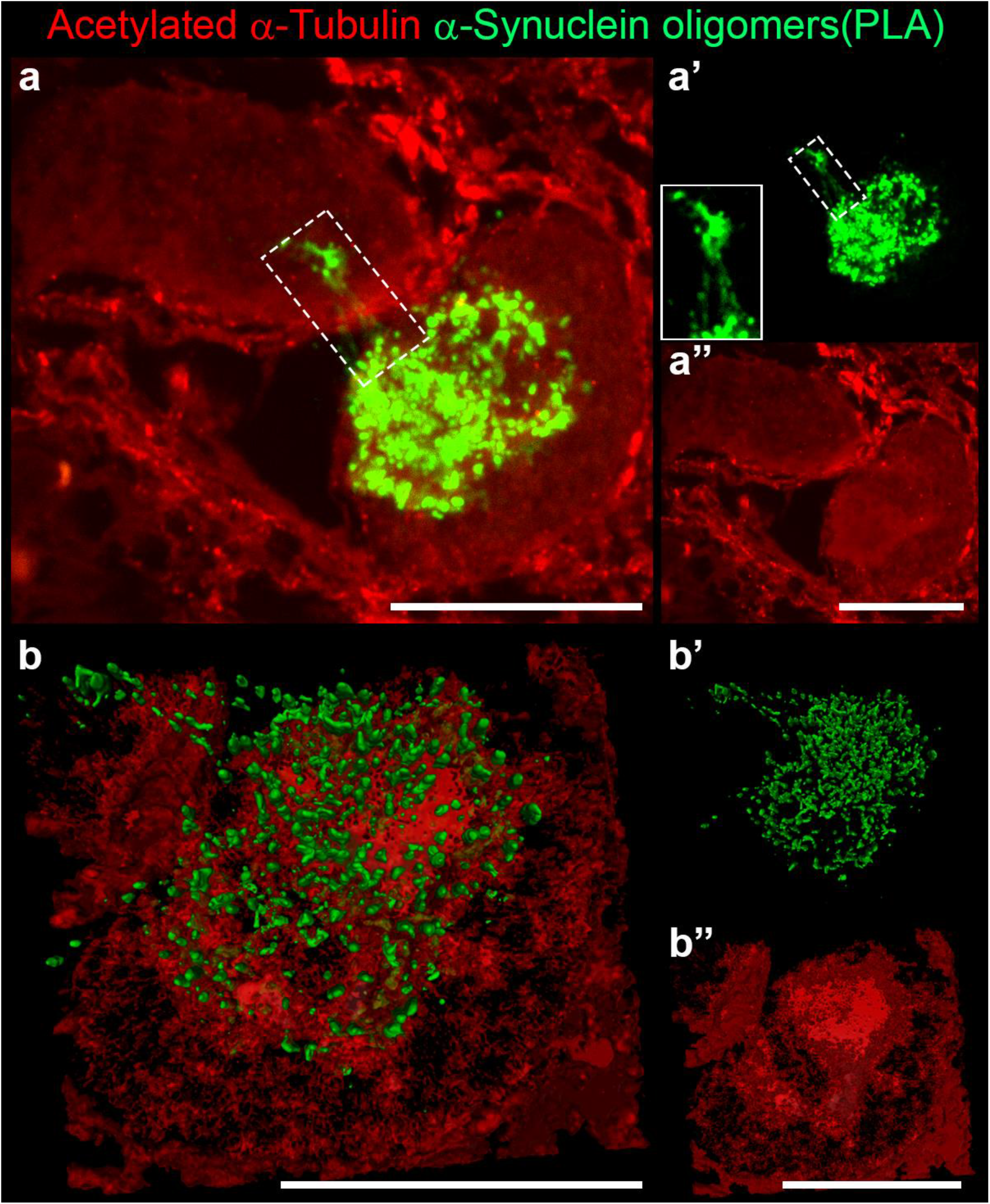
Tunnelling nanotube linking two neurons in *substantia nigra* of *post-mortem* human brain of PD patients. **a-a’’**α-Synuclein oligomers (**a-a’**) and acetylated α-Tubulin (**a’’**) mark the morphology of an undefined aggregate in neuronal cytoplasm. A tunnelling nanotube (white dashed rectangle, 2x magnified in **a’**) connects the two neurons and contains small oligomers. **b-b’’**3D reconstruction of the above confocal image with arivis Vision4D^®^ software. Scale bar, 10 μm.

## 5 Discussion

This study explores the morphogenesis of α-Synuclein aggregates, referred to as the histopathological hallmark of PD, with the goal to understand the role of α-Tubulin in triggering neurodegenerative processes. Importantly, we performed these analyses on human tissues obtained from PD patients aware that this is the only way to understand the pathological processes that experimental models may not be able to fully recapitulate. We disclose the redistribution of acetylated α-Tubulin in *post-mortem* human brain of PD patients and report its strong accumulation in neuronal cell bodies in the subcortical areas affected by LB pathology, from the dorsal motor nucleus of vagus to the *substantia nigra* ^61^. Notably, acetylated α-Tubulin decreases in the fibres of both the putamen and skin sweat glands suggesting that its rearrangement could play a role both in the central and peripheral nervous system. Hence, we linked the redistribution of acetylated α-Tubulin with α-Synuclein pathology and found that the accumulation of acetylated α-Tubulin in the cell soma precedes the formation of mature LBs, as revealed by the detailed analysis of the earliest forms of α-Synuclein aggregation i.e. oligomers. Starting from the staging of LB maturation that has very recently been described by Moors and colleagues ^47^, we applied the PLA approach thus unravelling a correlation between the acetylated form of α-Tubulin and oligomers, which supports further evidence to the emerging picture of LB morphogenesis, in which acetylated α-Tubulin may have an active part and play a driving-role.

To date, the evidence for a functional role of α-Tubulin acetylation is emerging in acute and toxic or gene-based models of parkinsonism^79^. An overall increase in α-Tubulin acetylation marks neuronal-like cells exposed to the neurotoxin MPP+^28^ and is detectable in dopaminergic fibres of the corpus striatum and in cell bodies of the *substantia nigra* dopaminergic neurons in mice injected with the neurotoxin MPTP ^30^. Similarly, neuroblastoma cells treated with 6-hydroxydopamine (6OHDA) display an increase in acetylated α-tubulin due to a reduction in the tubulin deacetylase enzyme SIRT2 ^80^. Interestingly, such an imbalance in the level of modified tubulin seems to precede the impairment of mitochondria transport in MPP+-treated cells ^28^. Thus, these studies converge on the hypothesis that increased acetylation of α-tubulin is implicated in the mechanisms of neurodegeneration. As regards genes linked to familial and non-familial forms of PD, including PARK2 (encoding for Parkin) and LRRK2 (encoding for Leucine-rich repeats kinase 2, LRRK2), the interplay between changes in acetylation of α-tubulin and functional defects in axonal transport is evident ^49,81^. Beyond the known role of Parkin in the stabilization of microtubules ^82,83^, it has been reported that in the PARK2 KO mouse model, acetylated α-tubulin has enhanced in dopaminergic neurons and fibres, and precedes the impairment of axonal mitochondria transport, as revealed by the clustering of mitochondria along the axons ^84^. To note, in dopaminergic neurons obtained from iPSCs, derived from patients carrying PARK2 mutations, acetylated α-tubulin is altered and displays a discontinuous and gapped staining on microtubules suggesting their overall weakness and dysfunction ^84^. Furthermore, primary fibroblasts obtained from skin biopsies carrying the LRRK2 mutation (p.G2019S) display an increase in acetylated α-tubulin in their cellular bodies ^85^. In contrast, Godena and colleagues demonstrated that LRRK2 pathogenic forms induce a decrease in acetylated α-tubulin, impairing and disrupting mitochondrial transport in cortical neurons and motor deficits in D. melanogaster, which are rescued increasing the acetylation on microtubules ^86^. Further to these debated results obtained *in vitro* and in animal models, our findings demonstrate, for the first time, that a redistribution of acetylated α-Tubulin occurs in patients affected with idiopathic PD and is detectable precisely in those brain regions that, at different stages of the pathology, are involved in α-Synuclein aggregation. Thus, our data shed light on the role of acetylated α-Tubulin as a reliable player in neurodegeneration processes underlying PD. Our study first demonstrates that neurons containing mature LBs are rarely positive for acetylated α-Tubulin. This could be due to the time-course of the events, in which the enrichment of acetylated α-Tubulin occurs in an early temporal window compared to the appearance of the final aggregated form of α-Synuclein. Our analysis of α-Synuclein oligomers, which are the earliest and most toxic species of α-Synuclein aggregates ^37,67,87^, allowed us to explore in detail the morphogenesis of LBs and propose a novel model of staging in which acetylated α-Tubulin and α-Synuclein oligomers cooperate (Fig. 7b). To date, in the sequence of events that leads to LB formation, α-Synuclein is first diffuse in the cytoplasm, gradually accumulates into irregularly-shaped aggregates, and, finally, into well-defined round-shaped inclusions such as pale bodies and LBs ^45^,^88^,^89^. Very recently, Moors and colleagues confirmed this staging and revealed details as to the framework of neurofilaments, ?-Tubulin, and phosphorylated Synuclein inside the aggregates using STED microscopy ^47^. Our study combined the PLA approach to detect α-Synuclein oligomers and immunofluorescence for acetylated α-Tubulin with spinning disk confocal microscopy, SoRa super-resolution, and 3D-reconstruction, to propose a novel multistage model for LB morphogenesis. This model involves acetylated α-Tubulin starting from stage 1, where it accumulates in the neuronal cell bodies prior to the appearance of α-Synuclein oligomers, to stage 6, where oligomers are localised mainly at the periphery of the inclusion and staining for acetylated α-Tubulin is weak (Fig.7b). The initial increase in α-Tubulin acetylation could be explained by defects in the activity of the enzymes that regulate acetylation on Lys40 of α-Tubulin, α-TAT1 ^90,91^ or its deacetylation, HDAC6 ^92,93^. However, we observed the decrease of acetylatedα-Tubulin in the cytoplasm during LB morphogenesis. We hypothesise that α-Synuclein might bind and sequester acetylated α-Tubulin thus reducing its level in the cytoplasm or, on the contrary, that accumulating acetylated α-Tubulin drives α-Synuclein oligomerization and aggregation into LBs. Our data showing the correlation between the volume of PLA puncta (i.e. α-Synuclein oligomers), and acetylated α-Tubulin (Fig. 9) point to the driving role of acetylated α-Tubulin. In stage 6, acetylated α-Tubulin is reduced inside ring-shaped aggregates (Fig. 7a). This could be related to the presence of the active phosphorylated form of HDAC6 which has recently been found to localise into mature α-Synuclein aggregates in *post-mortem* brain of PD patients ^58^. The question remains whether the increase of acetylated α-Tubulin during LB morphogenesis is protective or detrimental for neurons. The inhibition of HDAC6 with Tubastatin A in a rat model of PD-like neurodegeneration increases acetylated α-Tubulin and protects dopaminergic neurons reducing not only α-Synuclein toxicity and its phosphorylation at Ser129 but also neuroinflammation ^94^. In addition, Outeiro and colleagues revealed that the genetic and pharmacological inhibition of the other deacetylase of acetylated α-tubulin, SIRT2 ^93^, increases acetylated α-tubulin in cell models and reduces α-synuclein aggregation and toxicity^95^. Finally, studies on PD cybrid cells revealed that davunetide (NAP), a peptide promoting microtubule assembly, is able to reduce α-Synuclein oligomerization and restore microtubule-dependent trafficking by increasing the levels of acetylated α-Tubulin^96^. Hence, although data from experimental models suggest that acetylated α-tubulin control could be a promising strategy to reduce α-Synuclein aggregation and ameliorate PD conditions, its mechanism, namely reduction or stimulation, remains a matter of debate.

Interestingly, we report for the first time in *post-mortem* human brains the presence of threadlike structures that contain α-Synuclein oligomers and resemble tunnelling nanotubes. Until now, these structures have been seen exclusively in *in vitro* models. Tunnelling nanotubes were visualised between a donor and an acceptor neuron in co-cultures of mouse neuronal-like cells and were composed ofα-Synuclein fibrils in lysosomal vesicles that can seed α-Synuclein aggregation in the acceptor cell ^40^. More recently, another study elucidated the mechanism of α-Synuclein spreading with tunnelling nanotubes showing that α-Synuclein is able to modify the function of lysosomes and use these organelles to propagate and spread between cells ^97^. Moreover, Valdinocci and colleagues pointed out that α-Synuclein can bind migrating mitochondria within tunnelling nanotubes to increase its aggregation in mono and co-cultures of macrophages and neuroblastoma cells ^98^. Our study provides evidence that tunnelling nanotubes could be a strategy used by neurons in the central nervous system for the spreading of α-Synuclein pathology and, also interestingly, that acetylated α-Tubulin could be involved in the tunnelling nanotube formation. This hypothesis is based on the observation that the neurons associated with the threadlike structure containing α-Synuclein oligomers have increased acetylated α-Tubulin.

The neuronal imbalance of acetylated α-Tubulin is not exclusive to PD, but has been demonstrated in many neurodegenerative diseases^99^. In detail, changes in acetylated α-Tubulin have been reported in experimental models of Charcot-Marie-Tooth disease ^100 101^, Amyotrophic Lateral Sclerosis ^102,103^ Huntington’s ^104^ and Alzheimer’s disease ^105^. As regards *post-mortem* human brain, an imbalance of acetylated α-Tubulin is recorded in patients affected by neurodegenerative disease. In *post-mortem* striatal samples of Huntington’s disease (grade 3/4), Western blotting analysis revealed a dramatic reduction in acetylated α-Tubulin compared to controls, while total α-Tubulin levels were unaffected ^104^. A recent study on *post-mortem* human brain of Alzheimer’s disease patients at Braak stage III-IV reports a decrease in the level of acetylated α-Tubulin in the cortex and hippocampus ^31^. Interestingly, a previous work reported both a decrease in acetylated α-Tubulin and a reduction of total α-Tubulin in the hippocampus of Alzheimer’s disease patients compared to controls, with a net increase in the acetylation status of α-Tubulin ^106^. In the same work, Zhang and colleagues reported the accumulation of acetylated α-Tubulin in cell bodies of hippocampal neurons and this is similar to what we observed in PD brain and have described in our study. To date, the only evidence that the human brain in PD experiences an imbalance of acetylated α-Tubulin lies on biochemical analyses performed on the *substantia nigra* pars compacta from *post-mortem* brain of PD patients at Braak stage 5-6 ^31^. Our study investigates in detail its distribution in all brain areas affected by LB pathology as well as its subcellular distribution inside neurons. Our data usefully integrate the unique and aforementioned results obtained in PD ^31^ and, more importantly, open the way to investigate how the redistribution of this modified form of tubulin, which is crucial for the maintenance of the proper mechanical properties of microtubules, could impact the susceptibility of specific anatomical areas to neurodegeneration in PD and beyond.

In conclusion, this study reveals the contribution of acetylated α-Tubulin to LB morphogenesis in *post-mortem* human brains affected by PD. The completely unexplored link between the redistribution of acetylated α-Tubulin and α-Synuclein oligomerization in the human brain definitively supplies a novel perspective for investigating the drivers of pathological aggregation in PD.

## Declarations

## ACKNOWLEDGEMENTS

The authors thank all patients and families for their contribution, “Fondazione Grigioni per il Morbo di Parkinson” (Milan-Italy) for SM and FG salary and for long-lasting support to GC. We are grateful to Daniele Cartelli (Fondazione IRCCS Istituto Neurologico Carlo Besta, Milan, Italy) for critical reading and stimulating discussion and to Karen Doyle for reading and editing the paper. Part of this work was carried out at UNITECH NOLIMITS, an advanced imaging facility established by the Università degli Studi di Milano.

## AUTHOR CONTRIBUTIONS

All authors contributed to the study conception. MB, GG, SM, FG processed human samples. FG, GlG and HBI performed immunohistochemistry and multiple labelling experiments. SM, FG, HBI, MA performed confocal and super resolution imaging, as well as processing and analysis of images, also using arivis Vision 4D software. SM, GlG, AMC, FC performed pre-absorption experiments. SM, FG, GlG, HBI, AMC, FC, CR, GC performed data analysis. GC, GP, SM, FG, GG, IUI, AMC and CR designed research, analysed and interpreted the data, and contributed to writing the manuscript.

## COMPETING INTERESTS

The authors declare they have no conflict of interest.

## ETHICAL APPROVAL

The study was conducted according to the guidelines of the Declaration of Helsinki and approved by the Ethics Committee of the University of Milan (protocol code 66/21, 15 June 2021).

## ADDITIONAL INFORMATION

### Supplementary Figures 1-7

Supplementary Figure 1: Acetylated α-Tubulin antibody (mouse monoclonal 6-11-B) control.

Supplementary Figure 2: Acetylated α-Tubulin distribution in glial cells in *substantia nigra* of control subjects (Ctrl) and PD patients (PD).

Supplementary Figure 3: Acetylated α-Tubulin distribution in human cortex.

Supplementary Figure 4: Acetylated α-Tubulin distribution in white matter of frontal cortex in *post-mortem* human brain.

Supplementary Figure 5: α-Synuclein and Ser129P α-Synuclein distribution in α-Synuclein aggregates of *substantia nigra* in *post-mortem* human brain of PD patients.

Supplementary Figure 6: Histogram of the frequencies of PLA puncta volume in undefined (red) and ring-shaped (blue) aggregates.

Supplementary Figure 7: Tunnelling nanotubes in *substantia nigra* of PD patients.

### Supplementary Tables 1-3

Table 1. Demographic and clinical characteristics of the subjects included in the present study.

Table 2. Demographic and clinical characteristics of the skin biopsies included in the present study.

Table 3. Primary, secondary antibodies and kits used in this study.

### Supplementary movies

Movie 1: The movie refers to arivis 4D software 3D reconstruction of figure 6a’-a’’’.

Movie 2: The movie refers to arivis 4D software 3D reconstruction of figure 6b’-b’’’.

Movie 3: The movie refers to arivis 4D software 3D reconstruction of figure 6c’-c’’’.

Movie 4: The movie refers to arivis 4D software 3D reconstruction of figure 6d’-d’’’.

## Notes

### Competing Interest Statement

The authors have declared no competing interest.

