## Supplementary material for "Linking acetylated α-Tubulin redistribution to α-Synuclein pathology in brain of Parkinson’s disease patients": additional information.docx

^§^ co-last

^#^ **Correspondence:**

Graziella Cappelletti

Samanta Mazzetti

Supplementary Figures 1-7

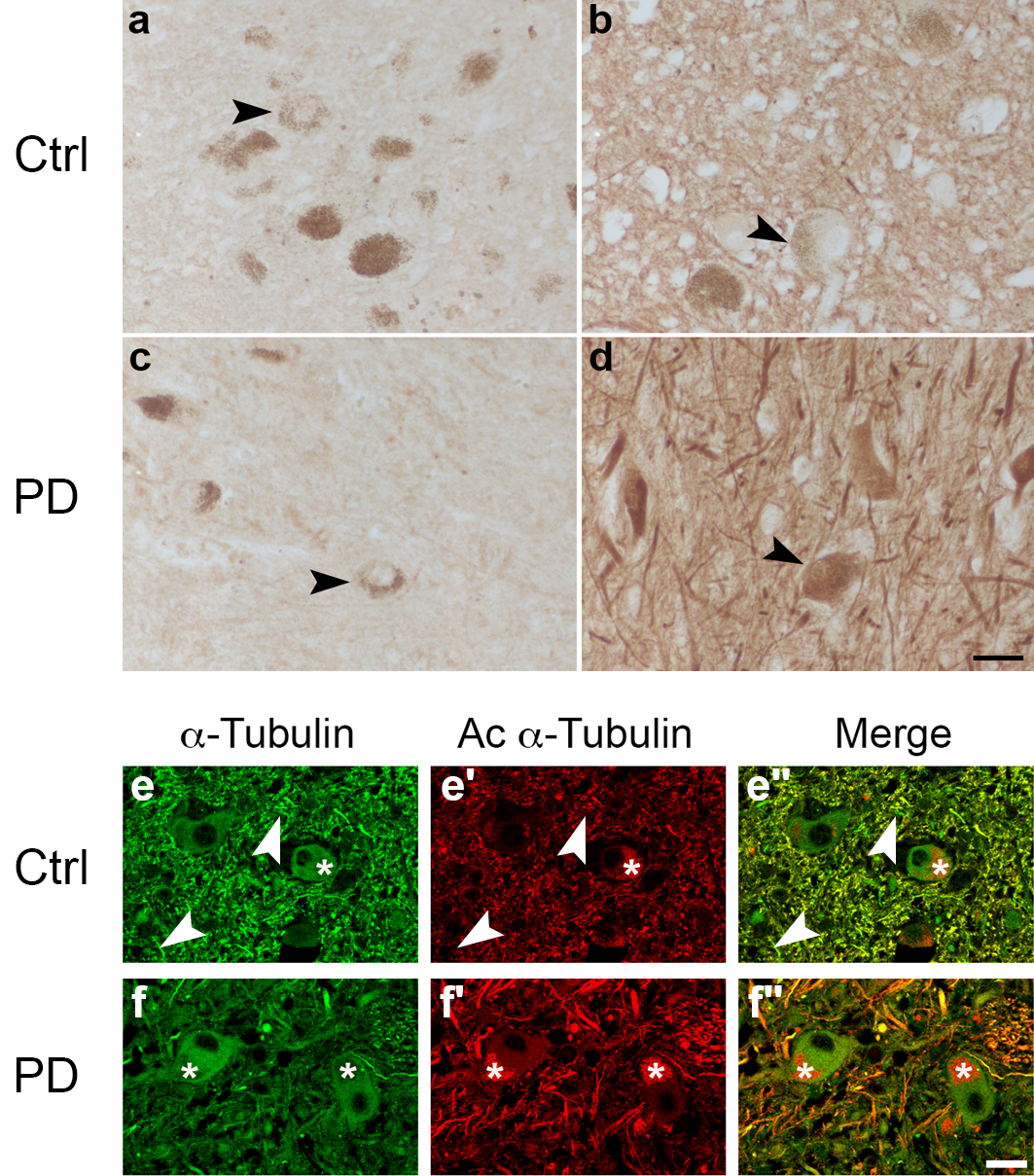

**Supplementary Figure 1.** Check for the specificity of acetylated α-Tubulin antibody (mouse monoclonal 6-11-B) in *post-mortem* human brain. (**a-d**): Sections containing *substantia nigra* of control subjects (Ctrl) and patients affected with Parkinson’s disease (PD) were incubated with anti-acetylated α-Tubulin pre-absorbed with tubulin purified from bovine brain and containing acetylated α-Tubulin (a, c), or with not-pre-absorbed anti-acetylated α-Tubulin (b, d). Staining for acetylated α-Tubulin is not detectable in sections incubated with the pre-adsorbed antibody. Dark brown signal is neuromelanin (black arrowheads). Scale bar, 40 μm. (**e-f”**): Sections containing dorsal motor nucleus of vagus of control subjects (Ctrl) and patients affected with Parkinson’s disease (PD) were double immunostained for total α-Tubulin (green) and acetylated α-Tubulin (red). As expected, neuronal cell bodies are positive for total α-Tubulin in controls (e, e’’) and PD patients (f, f’’). On the contrary, the staining for acetylated α-Tubulin in the neuronal cell bodies is evident exclusively in PD patients (f’, f’’) whereas cell bodies of controls are negative (e’, e’’). Furthermore, the specificity of the anti-acetylated α-Tubulin antibody that recognizes a fraction of the total α-Tubulin is confirmed by the staining of the neuropilar compartment, where some elements are exclusively positive for α-Tubulin (white arrowheads in e, e’, e’’). Asterisks indicates the nonspecific staining due to the presence of lipofuscin. Scale bar, 25 μm.

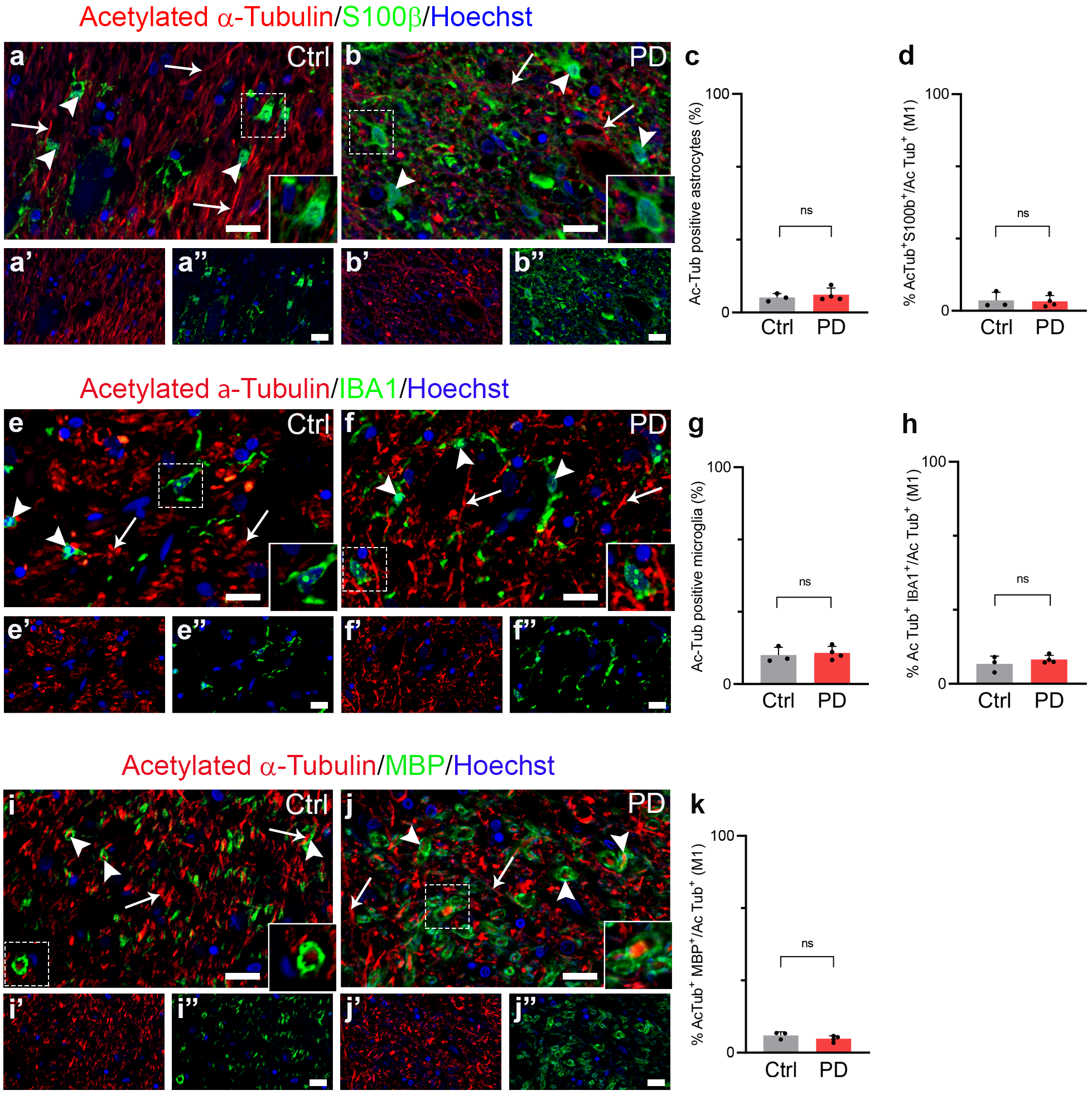

**Supplementary Figure 2.** Acetylated α-Tubulin distribution in glial cells in substantia nigra of control subjects (Ctrl) and PD patients (PD). Acetylated α-Tubulin is mainly localised in neuropil (white arrows) but not in astrocyte cell bodies (S100b, **a-b’’**; white arrowheads), microglia cell bodies (IBA1, **e-f’’**; white arrowheads), and MBP-positive oligodendrocytes (i-j’’; white arrowheads) in both control and PD samples. Insets: 1.5x magnified. Nuclei are counterstained with Hoechst. Graphs showing the quantitative analyses performed on astrocytes (**c-d**), microglial (**g-h**) and oligodendrocytes (**k**), report the percentage of glial cells positive for acetylated α-Tubulin (**c**: Ctrl, N = 3, 128 astrocytes vs PD, N = 4, 173 astrocytes; **g**: Ctrl, N = 3, 94 microglial cells vs PD, N = 4, 161 microglial cells) and the percentage of co-localisation between acetylated α-Tubulin and glial cells, expressed by Mander’s coefficient (M1) (**d, h, k**). Mann-Whitney test, ns. Scale bar, 20 μm.

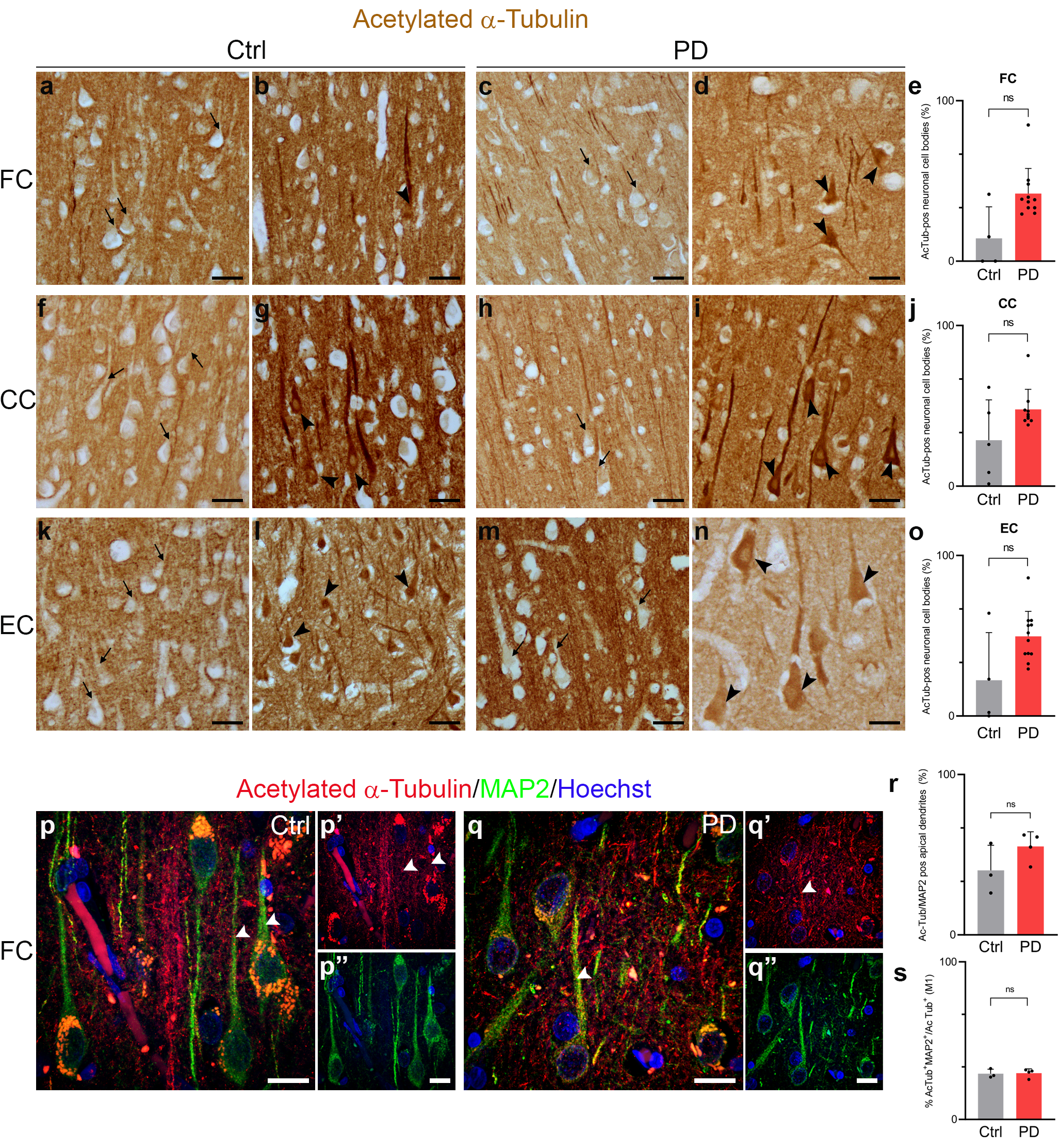

**Supplementary Figure 3.** Acetylated α-Tubulin distribution in human cortex. **a-o** Both control (**a, f, k**) and PD samples (**c, h, m**) show a homogeneous localisation of acetylated α-Tubulin inside apical dendrites (black arrows) in all three cortical regions. However, strong staining is present in the cell bodies of some pyramidal neurons (black arrowheads) in both control (**b, g, l**) and PD samples (**d, i, n**). Graphs show the percentage of acetylated α-Tubulin positive neuronal cell bodies (**e**: FC Ctrl, N = 4, 513 neurons vs PD, N = 11, 2212 neurons; **j**: CC Ctrl, N = 5, 672 neurons vs PD, N = 10, 1973 neurons. **o**: EC Ctrl, N = 4, 567 neurons vs PD, N = 12, 2016 neurons.). Scale bar, 40 μm. Mann-Whitney test, ns. **p-s** MAP2 stains the pyramidal neurons both in control (**p, p’’**) and PD (**q, q’’**) samples. Acetylated α-Tubulin is present is some apical dendrites in controls (white arrowheads; p, p’) while in PD it is also present in the soma of neurons (**q, q’**). Graphs show the percentage of apical dendrites positive for acetylated α-Tubulin (**r**; Ctrl, N = 3, 284 apical dendrites vs PD, N = 4, 428 apical dendrites) and the co-localisation between acetylated α-Tubulin and MAP2 (Mander’s coefficient, M1; **s**) Nuclei are stained with Hoechst. Scale bar, 10 μm. Mann-Whitney test, ns. FC: frontal cortex; CC: cingulate cortex; EC: entorhinal cortex.

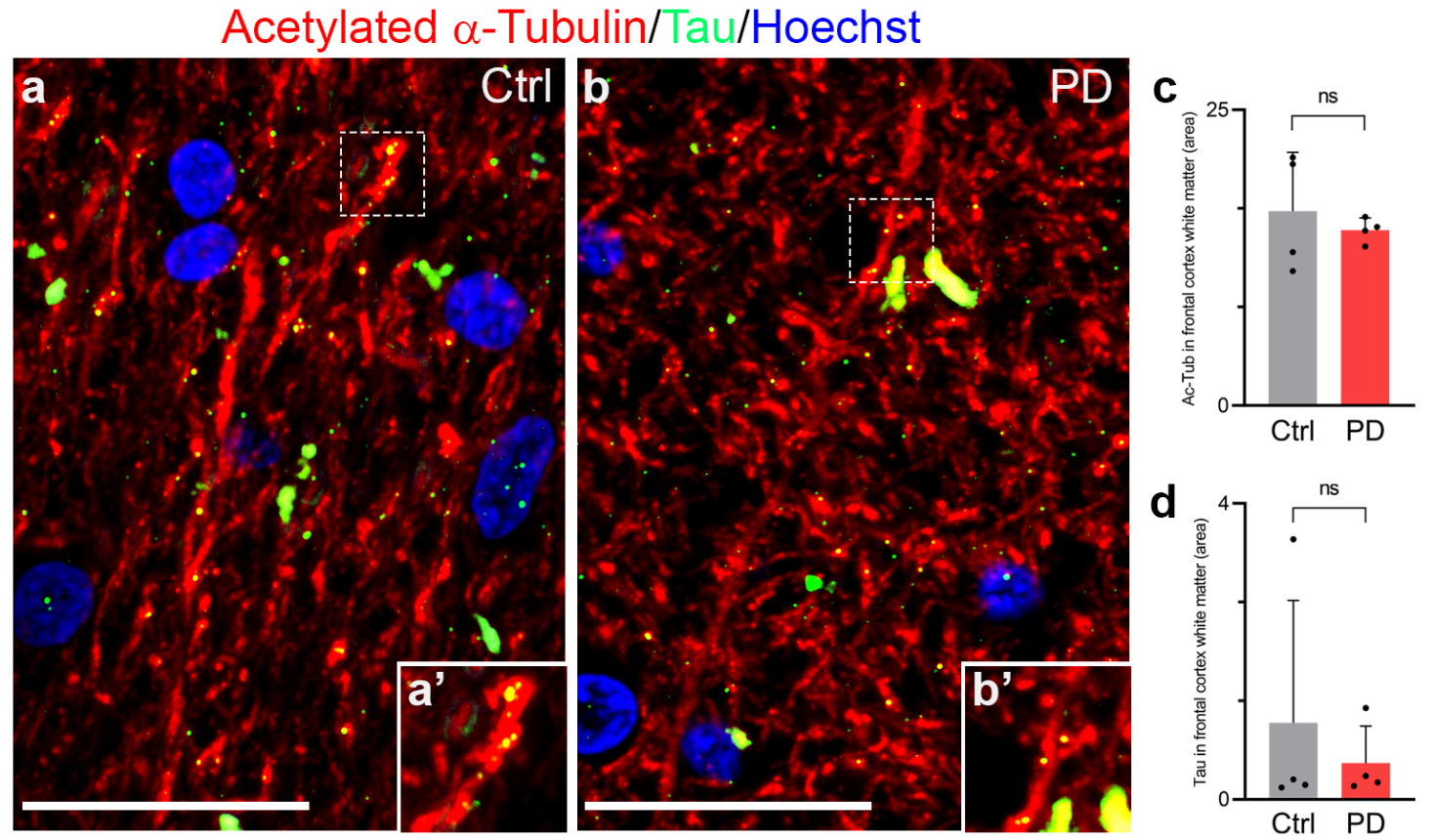

Tubulin is present is some apical dendrites

**Supplementary Figure 4**. Acetylated α-Tubulin distribution in white matter of frontal cortex in post-mortem human brain. Acetylated α-Tubulin stains axonal fibres of frontal cortex white matter both in control (**a-a’**) and PD (**b-b’**) samples. Tau displays a dotted staining and is distributed along the fibres both in controls (**a-a’**) and PD patients (**b-b’**). Inset: 2x magnified. Nuclei are counterstained with Hoechst. Graphs show the percentage of the area covered by acetylated α-Tubulin (**c**) and Tau (**d**) in axons. Mann-Whitney test, ns. Scale bar, 20 µm.

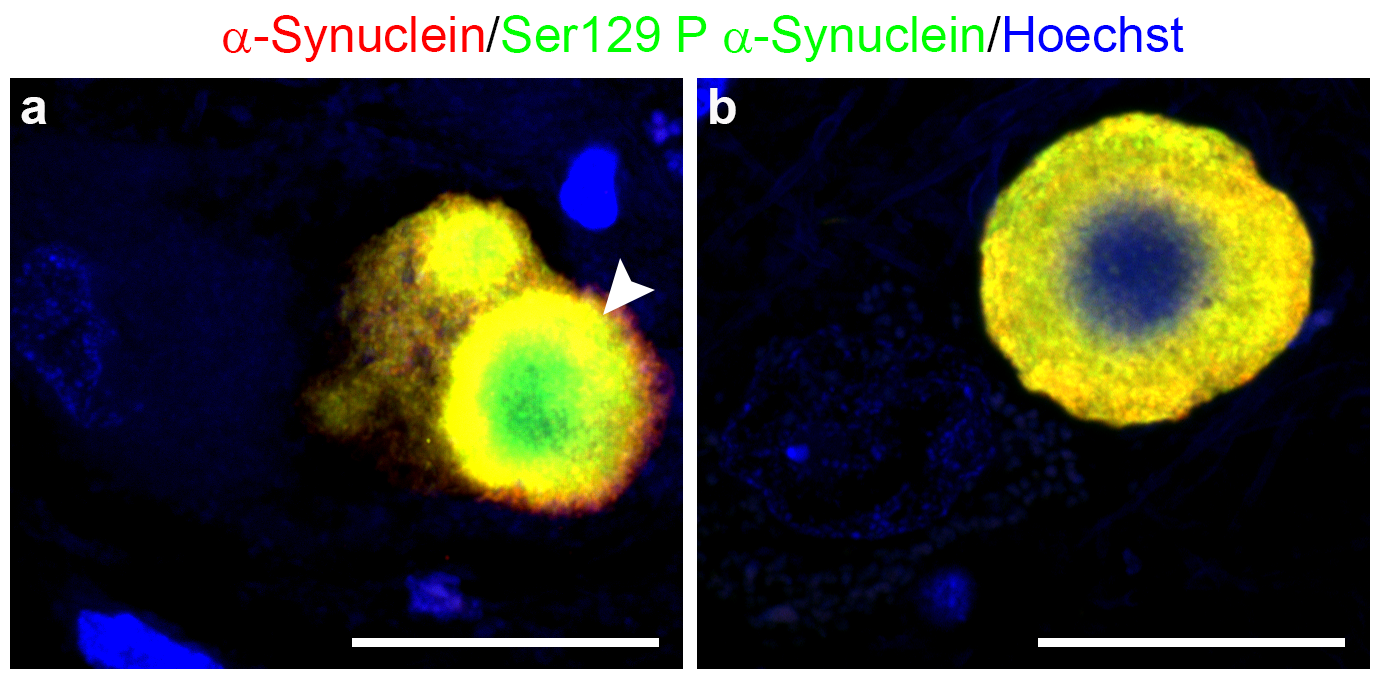

**Supplementary Figure 5**. α-Synuclein and Ser129P α-Synuclein distribution in α-Synuclein aggregates of substantia nigra in post-mortem human brain of PD patients. (**a**) Total and Ser129P α-Synuclein are inside aggregates without a defined shape. Both stainings form an external ring in which they co-localise, while Ser129P α-Synuclein is also present in the core region of the structure. In a mature LB (**b**), they co-localise in an external ring and Hoechst staining is detectable inside. Nuclei are counterstained with Hoechst. Scale bar, 20 μm.

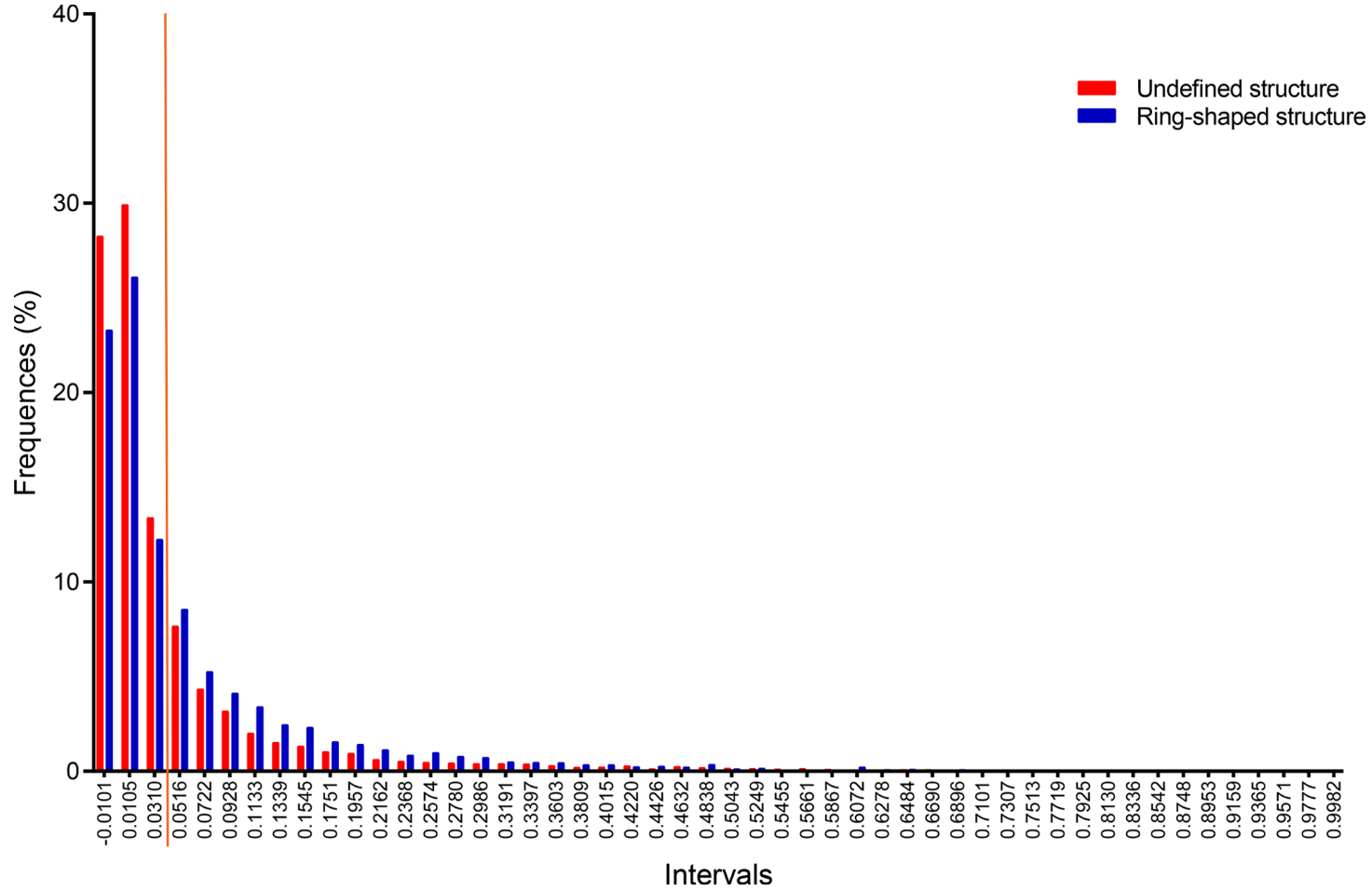
**Supplementary Figure 6**. Histogram of the frequencies of PLA puncta volume in undefined (red) and ring-shaped (blue) aggregates. At lower volume intervals, the frequency is higher for undefined aggregates compared to ring-shaped aggregates. The opposite is observed for PLA puncta larger than 0.0516 mm^3^. The orange line distinguishes the two intervals.

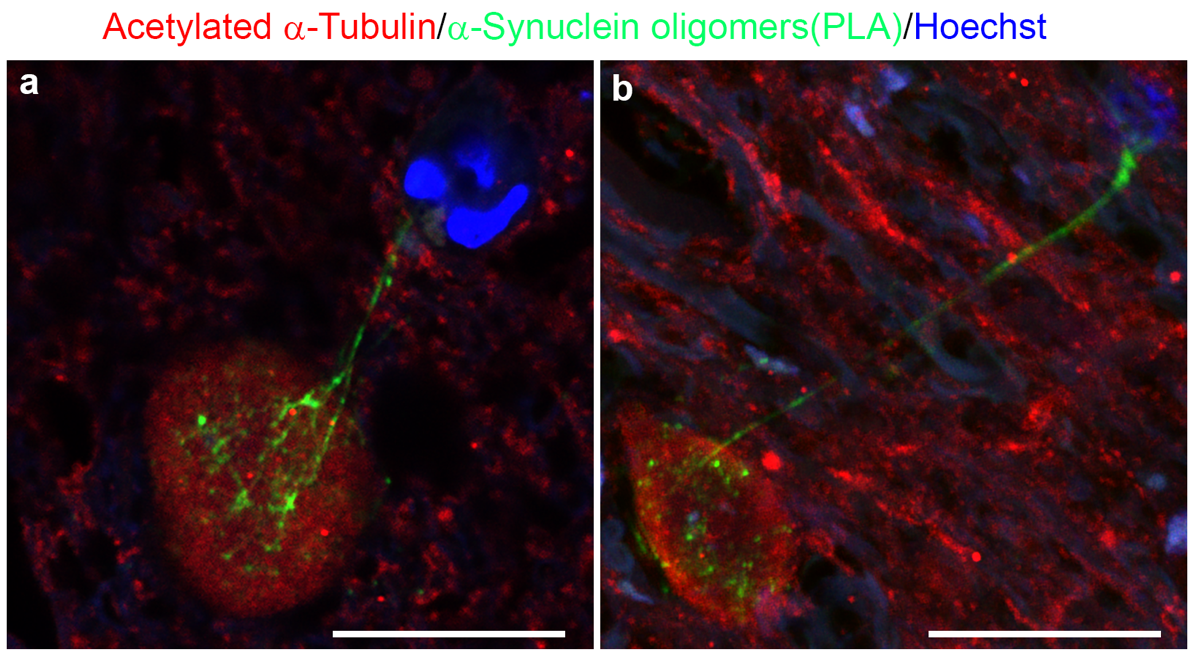

**Supplementary Figure 7**. Tunnelling nanotubes in substantia nigra of PD patients. α-Synuclein oligomers locate into threadlike structures that link a neuron accumulating acetylated α-Tubulin aggregates inside the soma to a vessel (**a**) and a glial cell (**b**). Scale bar, 25 μm.

Supplementary Tables 1-3

| Subject | Gender | Age at onset | Age at death | Disease duration (year) | Hoehn and Yahr |
| --- | --- | --- | --- | --- | --- |
| CTRL#1 | M | / | 71 | / | / |
| CTRL#2 | F | / | 93 | / | / |
| CTRL#3 | F | / | 82 | / | / |
| CTRL#4 | F | / | 64 | / | / |
| CTRL#5 | F | / | 91 | / | / |
| PD#1 | M | 40 | 59 | 19 | 5 |
| PD#2 | M | 57 | 71 | 14 | 5 |
| PD#3 | M | 57 | 75 | 18 | 3 |
| PD#4 | M | 59 | 75 | 16 | 3 |
| PD#5 | M | 59 | 87 | 28 | 3 |
| PD#6 | M | 62 | 73 | 11 | 3 |
| PD#7 | M | 62 | 80 | 18 | 4 |
| PD#8 | M | 72 | 84 | 12 | 4 |
| PD#9 | F | 43 | 72 | 29 | 5 |
| PD#10 | F | 53 | 91 | 38 | 5 |
| PD#11 | F | 59 | 79 | 20 | 4 |
| PD#12 | F | 65 | 84 | 19 | - |

**Supplementary Table 1**. Demographic and clinical characteristics of the subjects included in the present study.

| Subject | Gender | Age | Disease duration (year) |
| --- | --- | --- | --- |
| CTRL#1 | M | 82 | / |
| CTRL#2 | M | 40 | / |
| CTRL#3 | M | 66 | / |
| CTRL#4 | F | 58 | / |
| CTRL#5 | M | 62 | / |
| CTRL#6 | F | 61 | / |
| CTRL#7 | F | 57 | / |
| CTRL#8 | F | 61 | / |
| CTRL#9 | M | 39 | / |
| PD#1 | M | 41 | 6 |
| PD#2 | M | 56 | 8 |
| PD#3 | M | 61 | 9 |
| PD#4 | F | 31 | 7 |
| PD#5 | M | 48 | 8 |
| PD#6 | F | 69 | 24 |
| PD#7 | F | 51 | 14 |
| PD#8 | M | 54 | 1 |
| PD#9 | M | 52 | 21 |
| PD#10 | M | 71 | 1 |
| PD#11 | M | 64 | 7 |
| PD#12 | F | 78 | 9 |

**Supplementary Table 2.** Demographic and clinical characteristics of the skin biopsies included in the present study.

| Primary antibodies | | | |
| --- | --- | --- | --- |
| Antigen | Code | Host | Dilution |
| α-Synuclein | S3062 Merck | Rabbit | 1:2000 |
| Ser129 P α-Synuclein | ab51253Abcam | Rabbit | 1:1000 |
| Acetylated α-Tubulin | 6-11-B Sigma-Aldrich | Mouse | 1:1000 (IHC)  1:500 (brain IF)/1:4000 (skin IF) |
| Ionized calcium binding adapter molecule 1 (Iba1) | GTX 100042 GeneTex | Rabbit | 1:500 |
| Microtubule Associated Protein 2 (MAP2) | Ab5392 Abcam | Chicken | 1:500 |
| Mielin Basic Protein (MBP) | A0623 Dako | Rabbit | 1:1000 |
| Protein gene Product 9.5 (PGP 9.5) | AB1671-I Merck | Rabbit | 1:100 |
| S100β | 287006 Synaptic Systems | Chicken | 1:500 |
| Tau | 314012 Synaptic Systems | Rabbit | 1:200 |
| Tyrosine Hydroxylase (TH) | PA-18372 Thermo Fisher | Goat | 1:200 |
| Secondary antibodies | | | |
| Fluorochrome/Enzyme Antibody | Code | Host | Dilution |
| Alexa Fluor® 488 anti-goat | Jackson ImmunoResearch | Donkey | 1:600 |
| Alexa Fluor® 488 anti-mouse | AB150101 Abcam | Donkey | 1:200 |
| Alexa Fluor® 568 anti-mouse | A10037 Thermo Fisher | Donkey | 1:200 |
| Alexa Fluor® 488 anti-rabbit | A21206 Thermo Fisher | Donkey | 1:200 |
| Alexa Fluor® 647 anti-rabbit | A32795 Thermo Fisher | Donkey | 1:200 |
| Cy3 anti-chicken | Jackson ImmunoResearch | Donkey | 1:600 |
| EnVision System-HRP Labelled polymer anti-mouse | K4001 Dako Omnis | Goat | 1:1 |
| ImmPRESS™-AP anti- rabbit | MP-5401Vector | Horse | 1:1 |
| Commercial assay | | | |
| Duolink® in situ probe marker MINUS | DUO920101KT Merck | - | * |
| Duolink® in situ probe marker PLUS | DUO920091KT Merck | - | * |
| Duolink® In Situ Detection Reagents Red | DUO92008 Merck | - | * |
| EnVision FLEX DAB+SubstrateChromogen System | K3468 Dako | - | * |
| Fast Blue B salt | F3378-1G Merck | - | 1 mg/ml |
| Hoescht 33342 | 62249 Thermo Fisher | - | 1:5000 |

**Supplementary Table 3.** Primary, secondary antibodies and kits used in this study.

* used as indicated by the manufacture instruction.

Supplementary movies

**Movie 1.** The movie refers to arivis 4D software 3D reconstruction of figure 6a’-a’’’.

**Movie 2.** The movie refers to arivis 4D software 3D reconstruction of figure 6b’-b’’’.

**Movie 3.** The movie refers to arivis 4D software 3D reconstruction of figure 6c’-c’’’.

**Movie 4.** The movie refers to arivis 4D software 3D reconstruction of figure 6d’-d’’’.
